# Senotherapeutic potential against xeroderma pigmentosum

**DOI:** 10.1101/2025.05.22.655641

**Authors:** Xuebing Wang, Masako Nishida, Ai Yoshioka, Claire Yik-Lok Chung, Shinya Hashimoto, Hideki Tanizawa, Shinya Ohta, Ken-ichi Noma, Takeshi Fukumoto

## Abstract

Xeroderma pigmentosum (XP) is an inherited photoaging syndrome caused by mutations in genes involved in the nucleotide excision repair (NER) pathway. XP patients exhibit hypersensitivity to ultraviolet (UV) radiation, leading to accelerated skin aging and requiring lifelong sun avoidance. Here, we demonstrate that UV-induced DNA damage triggers cellular senescence and up-regulates senescence-associated secretory phenotype (SASP) genes in melanocytes derived from an XP patient. To explore the potential therapeutics for XP, we developed a cisplatin-based drug screening system and identified JAK inhibitors and curcuminoids as promising senomorphic agents. In addition, two classes of senolytic agents, BCL-2-like protein inhibitors and HSP90 inhibitors, effectively eliminate senescent melanocytes. Further analysis demonstrates that senomorphic treatment effectively counteracts senescence and reduces SASP gene expression in XP-derived melanocytes. Moreover, genes in senescence-related pathways, including the JAK/STAT, Type I interferon (IFN-I), and PI3K/AKT pathways, which are activated by both UV irradiation and cisplatin treatment, are down-regulated by senomorphic treatment. This study highlights a potential senotherapeutic strategy for XP, which may help alleviate photoaging symptoms in XP patients.

## Introduction

Xeroderma pigmentosum (XP) is an autosomal recessive disorder caused by mutations in the *XPA*−*XPG* genes, which encode proteins involved in the nucleotide excision repair (NER) pathway. These mutations lead to the classification of XP into subtypes, from XP-A to XP-G (Lehmann et al., 2011). In addition, the XP variant (XP-V) subtype is associated with a mutation in the *XPV* gene encoding DNA polymerase η, which is required for DNA replication through pyrimidine dimers (Lehmann *et al*., 2011). There are notable geographic differences in the incidence and subtype distribution of XP. In Western countries, XP occurs in approximately 2 to 4 cases per 1 million individuals, with XP-C being the most prevalent subtype (Kleijer et al., 2008; Robbins et al., 1974). In contrast, the incidence in Japan is significantly higher, ranging from 10 to 25 cases per 1 million individuals, which is approximately 10 times the rate observed in Western populations (Hirai et al., 2006). Notably, more than half of XP patients in Japan belong to the XP-A subtype, primarily due to a founder mutation in the *XPA* gene [g.15148G>C (c.390-1G>C)]. This mutation results in aberrant splicing of the *XPA* transcript, leading to negligible expression of the mutant gene (Imoto et al., 2013; Satokata et al., 1990).

XP patients exhibit extreme sensitivity to ultraviolet (UV) radiation and accelerated photoaging. Clinically, XP is characterized by the development of freckle-like pigmented or depigmented maculae and a heightened predisposition to skin cancers in sun-exposed areas (Lehmann *et al*., 2011; Moriwaki et al., 2017; Takemori et al., 2024). The underlying cause of XP is a deficiency in the NER pathway, which is responsible for repairing and removing UV-induced DNA photolesions such as cyclobutane pyrimidine dimers and 6-4 pyrimidine-pyrimidone photoproducts. In the absence of effective DNA repair, these lesions accumulate, leading to genomic mutations and a significantly increased risk of carcinogenesis (Schuch et al., 2017). This study focuses on the impact of UV-B exposure, specifically within the 280-315 nm wavelength range, which accounts for the majority of UV-induced DNA lesions in the epidermis (Schuch *et al*., 2017). XP patients, particularly those under the age of 20, have a significantly higher incidence of skin cancer. They exhibit a 2,000-fold increase in melanoma cases and a 10,000-fold increase in non-melanoma skin cancers compared to non-XP individuals (Bradford et al., 2011; Cleaver, 1968; Nishigori et al., 2019; Takemori *et al*., 2024). In addition to cutaneous manifestations, 20-55% of XP patients also exhibit neurological dysfunction due to central and peripheral neurodegeneration (Lehmann *et al*., 2011; Moriwaki *et al*., 2017). Despite the severity of XP symptoms, no radical cure exists. Symptomatic management, including sun protection and early detection and excision of skin cancer lesions, remains the primary clinical intervention (Moriwaki *et al*., 2017). Among the XP subtypes, XP-A patients exhibit the most severe cutaneous and neurological symptoms due to their severely impaired DNA repair capacity, highlighting the critical need for novel therapeutic strategies (Nishigori *et al*., 2019; Takemori *et al*., 2024).

XP results from defects in the NER pathway, leading to an accumulation of DNA damage following UV irradiation (Kamenisch and Berneburg, 2009). This DNA damage can trigger cellular senescence, a state of stable cell-cycle arrest (Munoz-Espin and Serrano, 2014). The premature aging symptoms observed in XP patients, particularly in the skin, may be attributed to accelerated senescence induced by UV exposure. Consequently, targeting cellular senescence represents a promising therapeutic approach for the treatment of XP. In recent years, accumulating evidence has linked cellular senescence to a variety of human diseases, establishing it as a potential therapeutic target (Di Micco et al., 2021; Wang et al., 2024a). Clinical interventions aimed at modulating senescence are referred to as senotherapy. Senotherapeutic agents are categorized into two groups: senolytic agents, which selectively eliminate senescent cells via apoptosis, and senomorphic agents, which inhibit the production of senescence-associated secretory phenotype (SASP) factors (Di Micco *et al*., 2021; Wang *et al*., 2024a). At present, the precise role of cellular senescence in the etiology of XP, and, more importantly, the potential effectiveness of senotherapeutic strategies in treating XP remains to be determined.

Senolytic agents, such as ABT-263 (inhibitor of BCL-2-like proteins), 17-DMAG (HSP90 inhibitor), and digoxin (Na^+^/K^+^ ATPase pump inhibitor), have been shown to eliminate senescent cells, highlighting the therapeutic potential for various diseases such as cancer, pulmonary fibrosis, and atherosclerosis (Fuhrmann-Stroissnigg et al., 2017; Triana-Martinez et al., 2019; Wang et al., 2024b). BCL-2-like proteins (e.g., BCL-2, BCL-W, BCL-XL, and MCL-1) are up-regulated in senescent cells and promote resistance to apoptosis. ABT-263 specifically inhibits these BCL-2-like proteins, thereby inducing apoptosis in senescent cells (Chang et al., 2016). While the effects of senolytic agents are preferentially restricted to senescent cells, ABT-263 is known to cause platelet toxicity, raising safety concerns for its clinical application (Kirkland and Tchkonia, 2020; Schoenwaelder et al., 2011; Wilson et al., 2010).

Senomorphic agents suppress the production of SASP factors, thereby inhibiting the propagation of senescent cells by blocking both autocrine and paracrine senescence (Di Micco *et al*., 2021; Wang *et al*., 2024a). Notably, senomorphic agents such as metformin, rapamycin, and baricitinib target key SASP regulators, including NF-κB, mTOR, and JAK, respectively, leading to a reduction in SASP factor production (Herranz et al., 2015; Liu et al., 2019; Moiseeva et al., 2013). In preclinical models, these senomorphic agents have shown efficacy in treating various diseases, including certain cancers, age-related dysfunctions, and progeria syndromes (Arnold et al., 2021; Herranz *et al*., 2015; Liu *et al*., 2019; Moiseeva *et al*., 2013; Xu et al., 2015). Moreover, the natural product curcumin and its analog EF24 also exhibit senomorphic properties by suppressing NF-κB, suggesting their potential for treating various senescence-related diseases (Li et al., 2019; Yin et al., 2016; Zia et al., 2021). Collectively, these studies indicate that senotherapeutic strategies employing senolytic and senomorphic agents could be effective for aging-related syndromes, although their efficacy in XP remains undetermined. In this study, we identify effective senolytic and senomorphic agents and propose senomorphic agents, JAK inhibitors (baricitinib and upadacitinib) and curcuminoids (curcumin and EF24), as promising candidates for the treatment of XP.

## Results

### Senescence induction of XP-derived melanocytes (XP-iMCs) by UV irradiation

Since melanocytes, rather than keratinocytes, are more prone to become senescent in aged skin (Victorelli et al., 2019; Waaijer et al., 2016), we hypothesized that melanocytes from XP patients would have an increased tendency to undergo senescence upon UV irradiation due to their deficiency in the NER pathway. This heightened susceptibility could contribute to accelerated skin aging observed in XP patients. To test this hypothesis, we examined the response of XP patient-derived melanocytes to UV irradiation. In our preparatory study, we established a system in which induced pluripotent stem cells (iPSCs) derived from an XP-A patient and a healthy control individual were differentiated into melanocytes, referred to as XP-iMCs and HC-iMCs, respectively (Takemori *et al*., 2024). We employed these iPSC-derived melanocytes after verifying their physiological and functional identity, as evidenced by their characteristic morphology and the expression of the melanocyte markers MITF and SOX10 (Hosaka et al., 2019; Takemori *et al*., 2024) (**Supplementary Figure 1**).

To investigate UV-induced senescence of XP-iMCs, we irradiated XP-iMCs and HC-iMCs (as a control) with 150 J/m^2^ UV-B and assessed senescence markers one day after irradiation (**Figure 1a**). This UV dose was chosen based on our previous study, which showed that 150 J/m^2^ UV-B inhibits cell proliferation and induces the expression of cytokines, including potential SASP factors (Fukumoto et al., 2016; Fukumoto et al., 2025; Takemori *et al*., 2024). We first performed immunofluorescence staining for the DNA damage marker γH2AX and observed stronger γH2AX staining in UV-irradiated XP-iMCs compared to unirradiated cells (**Figure 1b**). In contrast, HC-iMCs showed minimal UV-dependent increase in γH2AX staining. These results indicate that UV-induced DNA lesions are not efficiently repaired in XP-iMCs due to their NER deficiency, leading to the accumulation of DNA damage.

**Figure 1.**
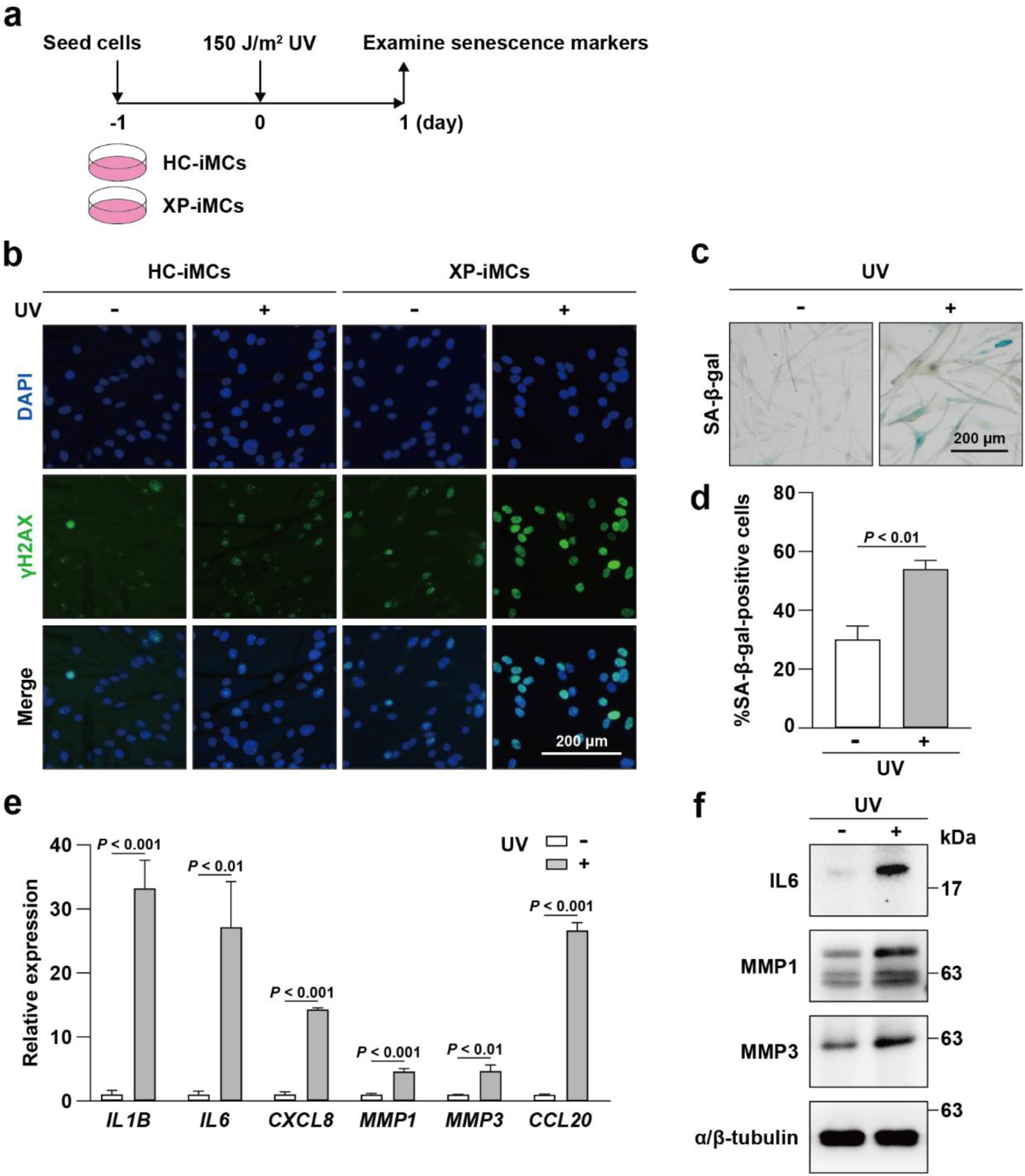
Senescence induction of XP-iMCs by UV irradiation. **(a)** Schematic representation of the UV irradiation procedure. HC-iMCs and XP-iMCs are melanocytes differentiated from HC-iPSCs and XP-iPSCs, respectively, derived from a healthy control individual and xeroderma pigmentosum patient (**Materials and Methods**). Cells were analyzed one day after exposure to 150 J/m^2^ UV, as shown in **panels b-f**. **(b)** Effect of UV irradiation on HC-iMCs and XP-iMCs. DNA damage marker γH2AX (green) was visualized by immunofluorescence microscopy and is shown alongside DAPI-stained nuclei (blue). **(c)** Senescence-associated β-galactosidase (SA-β-gal) staining of XP-iMCs with or without UV irradiation. **(d)** Quantification of SA-β-gal-positive XP-iMCs irradiated with UV. SA-β-gal staining images, as exemplified in **panel c**, were used for analysis. **(e)** Up-regulation of SASP genes in XP-iMCs irradiated with UV. RT-qPCR was performed to assess the expression of the indicated SASP genes. Expression levels in non-irradiated XP-iMCs were set to 1. **(f)** Western blot analysis of SASP factor expression in XP-iMCs with and without UV irradiation. α/β-tubulin serves as a loading control. In **panels d** and **e**, data are represented as means and standard deviations (n=3). Statistical significance was assessed using a two-sided Student’s *t*-test.

We next investigated whether accumulated DNA damage in XP-iMCs triggers cellular senescence. To do so, we monitored senescence-associated β-galactosidase (SA-β-gal) activity, a typical senescence marker, in XP-iMCs (Dimri et al., 1995). We observed that UV irradiation significantly increased the number of SA-β-gal-positive XP-iMCs (**Figure 1c, d**). In addition, we performed RT-qPCR and western blot analyses to assess the expression of SASP factors in XP-iMCs irradiated or unirradiated with UV. We found that both mRNAs and protein levels of certain SASP factors were more abundant in UV-irradiated XP-iMCs compared to unirradiated cells (**Figure 1e, f**). Taken together, these results indicate that DNA damage induced by UV irradiation accumulates in XP-iMCs and triggers cellular senescence, accompanied by the expression of SASP factors.

### Cisplatin sensitivity of XP-iMCs

As described above, UV irradiation induces cellular senescence of XP-iMCs, which may contribute to aging phenotypes in the skin of XP patients. Conversely, senotherapeutic strategies employing senolytic or senomorphic agents could offer novel clinical interventions for treating XP. To explore this possibility, we established a drug screening system using XP-iMCs. A key challenge in this screening process was the uneven effects of UV irradiation, as factors such as cell positioning in culture dishes may influence UV exposure. To overcome this issue, we considered using cisplatin treatment as an alternative to UV irradiation. Cisplatin-induced DNA lesions and damage can be precisely controlled by adjusting the cisplatin concentration in the culture medium, ensuring a uniform effect on all cells, regardless of their position in culture dishes. More importantly, cisplatin induces intrastrand DNA crosslinks, which are repaired by the NER pathway in the same manner as UV-induced pyrimidine dimmers (**Supplementary Figure 2**). Therefore, we propose that cisplatin treatment can effectively mimic UV irradiation as a senescence inducer in our drug screening system.

If cisplatin treatment and UV irradiation have similar effects on XP-iMCs, they should exhibit comparable sensitivity to both treatments. To test this possibility, we performed cell viability assays to examine the cisplatin sensitivity of XP-iMCs. For comparison, we included four cell lines: normal human epidermal melanocytes (NHEM), HC-iMCs, XP-iMCs, and XP-iMCs expressing wild-type XPA fused to Flag epitope (XP-iMCs + Flag-XPA). We observed that XP-iMCs were more sensitive to cisplatin than NHEM and HC-iMCs and that this cisplatin sensitivity was rescued by the expression of wild-type XPA in XP-iMCs (**Figure 2a**). Furthermore, we estimated the half-maximal inhibitory concentration (IC_50_) of cisplatin for these cell lines and found that the cisplatin IC_50_ value for XP-iMCs was significantly lower than those for other cell lines (**Figure 2b**). Additionally, we examined XPA expression levels in the respective cell lines using western blot and immunofluorescence (**Figure 2c, d**). As expected, a very strong XPA signal was detected in XP-iMCs expressing exogenous Flag-XPA, whereas endogenous XPA expression was observed at lower levels only in NHEM and HC-iMCs (**Figure 2c**). A previous study reported that the endogenous XPA is barely expressed in XP patients due to a splicing mutation in the *XPA* gene, which prevents the accumulation of functional *XPA* transcripts (Satokata *et al*., 1990). Consistent with this report, endogenous XPA was not detected in XP-iMCs. These results collectively suggest that XP-iMCs are hypersensitive to cisplatin due to the XPA depletion caused by the *XPA* mutation.

**Figure 2.**
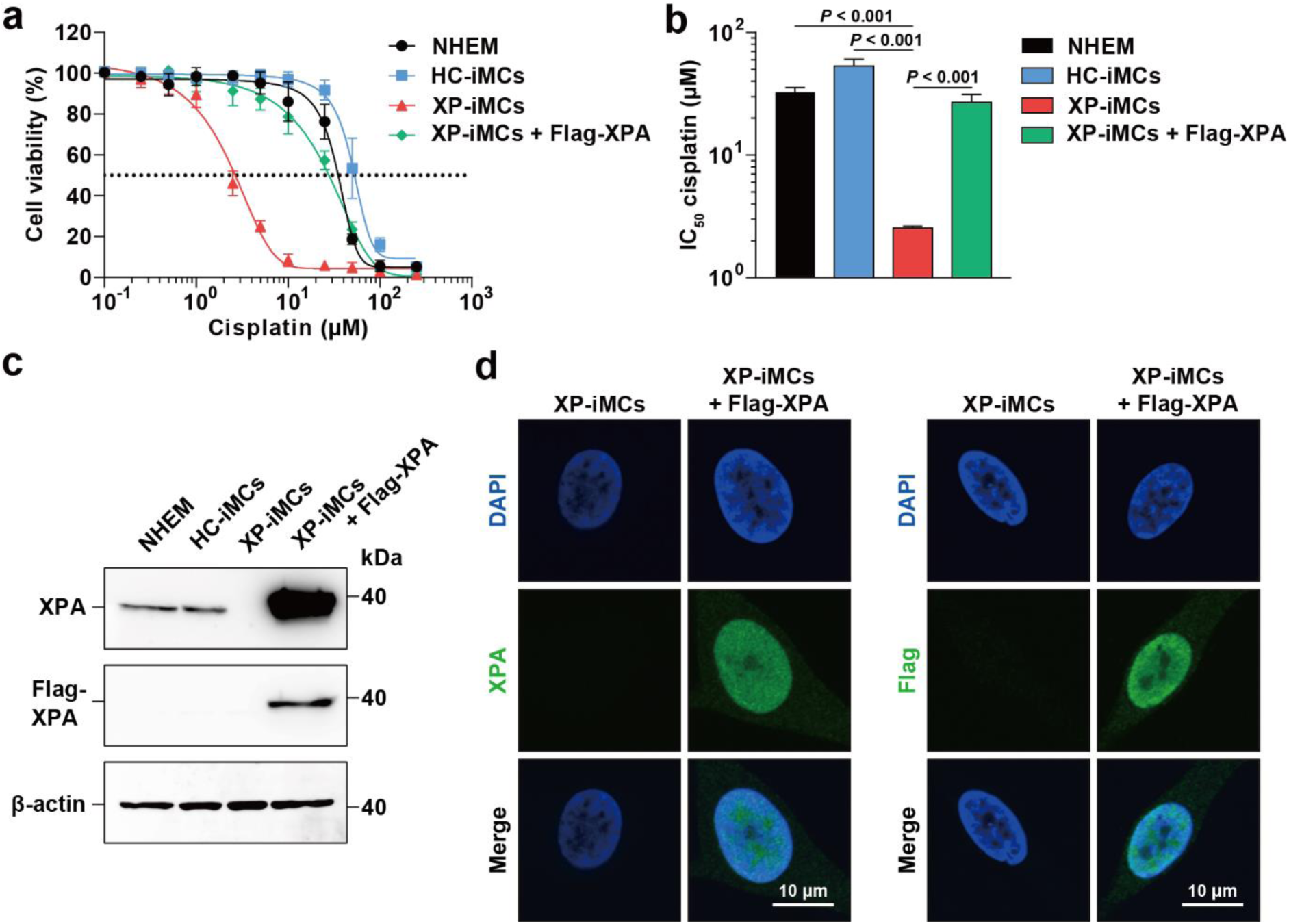
Expression of wild-type XPA suppresses cisplatin sensitivity of XP-iMCs. **(a)** Cisplatin dose-response curves for NHEM, HC-iMCs, XP-iMCs, and XP-iMCs expressing wild-type XPA tagged with a Flag epitope (Flag-XPA). Cell numbers were measured after 72 hours of culture at the indicated cisplatin concentrations. Data represent means ± standard deviations (n=3). **(b)** Cisplatin IC_50_ values for the indicated cell lines, estimated from the dose-response curves shown in **panel a**. Data represent means and standard deviations (n=3). Statistical significance was assessed using one-way ANOVA and Tukey’s test. **(c)** Western blot analysis of XPA protein expression in the indicated cell lines. XPA was detected using an anti-XPA antibody (top), and Flag-tagged XPA was detected using an anti-Flag M2 antibody (middle). β-actin serves as a loading control (bottom). **(d)** Immunofluorescence visualization of Flag-XPA proteins in XP-iMCs. XPA was detected using an anti-XPA antibody (left), and Flag-tagged XPA was detected using an anti-Flag M2 antibody (right).

### Senescence induction of XP-iMCs by cisplatin treatment

Since XP-iMCs are sensitive to cisplatin, we predicted that DNA damage derived from cisplatin-induced DNA lesions would trigger cellular senescence. To test this possibility, we investigated whether cisplatin treatment accumulates DNA damage and induces senescence of XP-iMCs. Briefly, XP-iMCs and HC-iMCs were seeded and cultured with a medium containing 1 μM cisplatin for 6 days, followed by the detection of DNA damage and senescence markers (**Figure 3a**). γH2AX-positive XP-iMCs were readily detected on days 4 and 6, suggesting that at least 4 days are required for XP-iMCs to accumulate detectable DNA damage under these conditions (**Figure 3b**). In contrast, γH2AX-positive cells were not detected for HC-iMCs, suggesting that the *XPA* mutation in XP-iMCs contributes to the observed accumulation of DNA damage caused by cisplatin. Notably, XP-iMCs, but not HC-iMCs, were also sensitive to UV irradiation (**Figure 1b**), suggesting that the enhanced sensitivity of XP-iMCs to both UV and cisplatin is consistently dependent upon the *XPA* mutation.

**Figure 3.**
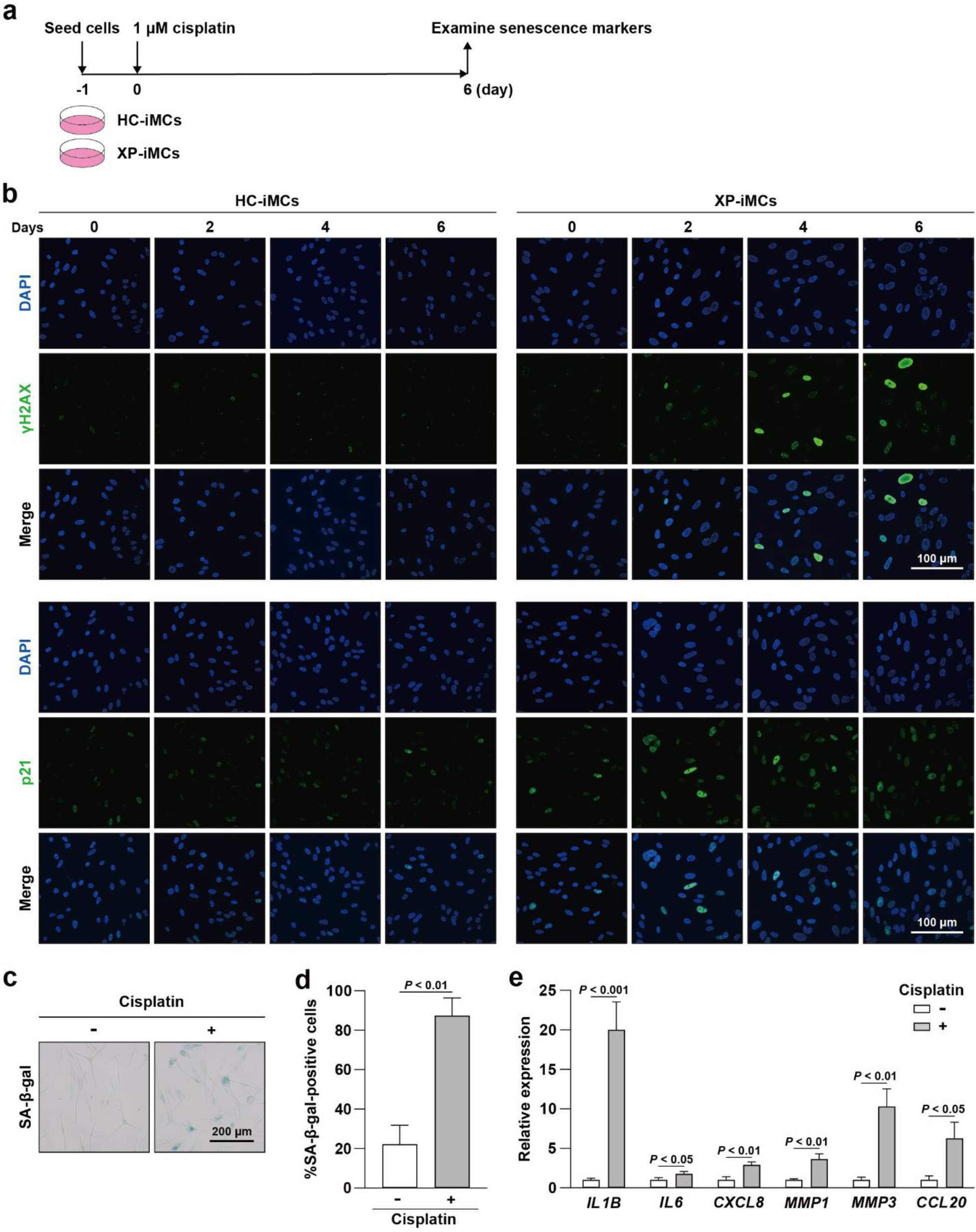
Cisplatin-induced senescence of XP-iMCs. **(a)** Schematic representation of the cisplatin treatment procedure. **(b)** Senescence induction by cisplatin treatment. HC-iMCs (left four columns) and XP-iMCs (right four columns) were treated with 1 μM cisplatin at the indicated durations (days) and subjected to immunofluorescence visualization of the senescence markers γH2AX (top three rows) and p21 (bottom three rows). **(c)** SA-β-gal staining of XP-iMCs treated or untreated with cisplatin. **(d)** Percentages of SA-β-gal-positive XP-iMCs treated or untreated with cisplatin. SA-β-gal-positive cells were quantified as described in **Supplementary Figure 3**. **(e)** Up-regulation of SASP genes in XP-iMCs treated with cisplatin. RT-qPCR analysis was performed on XP-iMCs treated or untreated with cisplatin to measure the expression levels of the indicated SASP genes. Expression levels in untreated XP-iMCs were set to 1. In **panels d** and **e**, data are represented as means and standard deviations (n=3). Statistical significance was assessed using a two-sided Student’s *t*-test.

We next examined the senescence induction of XP-iMCs by cisplatin treatment. To do this, we monitored the cell-cycle regulator p21, which plays a key role in senescence induction (Di Micco *et al*., 2021). p21 signals were enhanced in XP-iMCs from day 2, with most of cells becoming p21-positive at day 4 and 6. In parallel, γH2AX signals increased on day 4, implying that the undetectable level of DNA damage on day 2 was likely sufficient to induce senescence. Moreover, p21 signals were not increased by cisplatin treatment in HC-iMCs, suggesting that cisplatin-induced DNA damage drives senescence of XP-iMCs.

Since p21 is also up-regulated in quiescent cells (Di Micco *et al*., 2021), we assessed senescence more specifically by examining SA-β-gal activity. Consistently, SA-β-gal signals were readily detected in cisplatin-treated XP-iMCs (**Figure 3c**). To objectively quantify the senescence cell population, we employed SPiDER-βGal staining combined with flow cytometric analysis (Doura et al., 2016; Wang *et al*., 2024b) (**Supplementary Figure 3**). This analysis revealed a significant increase in the percentage of SA-β-gal-positive XP-iMCs treated with cisplatin compared to untreated cells (**Figure 3d**). Additionally, SASP genes were up-regulated in cisplatin-treated XP-iMCs (**Figure 3e**). Collectively, these results indicate that cisplatin treatment causes DNA damage and senescence of XP-iMCs. Furthermore, they suggest that cisplatin could serve as a senescence inducer for senotherapeutic drug screening.

### BCL-2-like protein inhibitors and HSP90 inhibitors as effective senolytic agents

Senotherapeutic agents can be categorized into senolytic agents, which selectively eliminate senescent cells, and senomorphic agents, which inhibit the production SASP factors (Di Micco *et al*., 2021). Our initial goal was to identify potential senolytic agents capable of eliminating cisplatin-induced senescent XP-iMCs. To this end, we reviewed recent literature and selected 12 senolytic agents based on their reported effects on senescent cells and mechanisms of action (Wang *et al*., 2024a) (**Supplementary Table 1**). In our initial drug screening approach, we simultaneously treated XP-iMCs with cisplatin and senolytic agents, expecting cisplatin to induce senescence, followed by selective elimination of senescent cells by the senolytic agents (**Supplementary Figure 4a**). Unexpectedly, increasing concentrations of senolytic agents impaired cell viability regardless of cisplatin treatment (**Supplementary Figure 4b**). We speculated that the lack of selectivity is due to the inherent vulnerability of melanocytes to senolytic agents. Supporting this notion, we observed that the viability of three melanocyte cell lines treated with ABT-263 was significantly lower than that of treated keratinocytes (**Supplementary Figure 4c**).

Based on these results, we predicted that a sequential strategy, in which senescence is first induced with cisplatin followed by senolytic treatment, may be more effective (**Figure 4a**). This approach is expected to take advantage of the heightened susceptibility of senescent melanocytes to senolytic agents compared to non-senescent melanocytes. Sequential treatment revealed that XP-iMCs were more sensitive to two categories of senolytic agents: BCL-2-like protein inhibitors (ABT-263, A-1331852, A-1155463, and S63845) and HSP90 inhibitors (17-DMAG and ganetespib) (**Figure 4b**). It has been shown that these HSP90 inhibitors exhibit senolytic effects on a cell line from a mouse model of a human progeroid syndrome, which is sensitive to DNA damage, supporting the validity of the screening system (Fuhrmann-Stroissnigg *et al*., 2017). Taken together, these results indicate the senolytic agents which target the BCL-2-like proteins or HSP90 selectively eliminate senescent XP-iMCs. Although certain senolytic agents appear effective in eliminating senescent XP-iMCs, melanocytes are more vulnerable to senolytic agents compared to keratinocytes. This raises concerns about the dose optimization of senolytic agents for XP treatment, specifically targeting senescent melanocytes while sparing non-senescent melanocytes. Therefore, identifying an alternative group of senotherapeutic candidates, such as senomorphic agents, could be beneficial for XP treatment.

**Figure 4.**
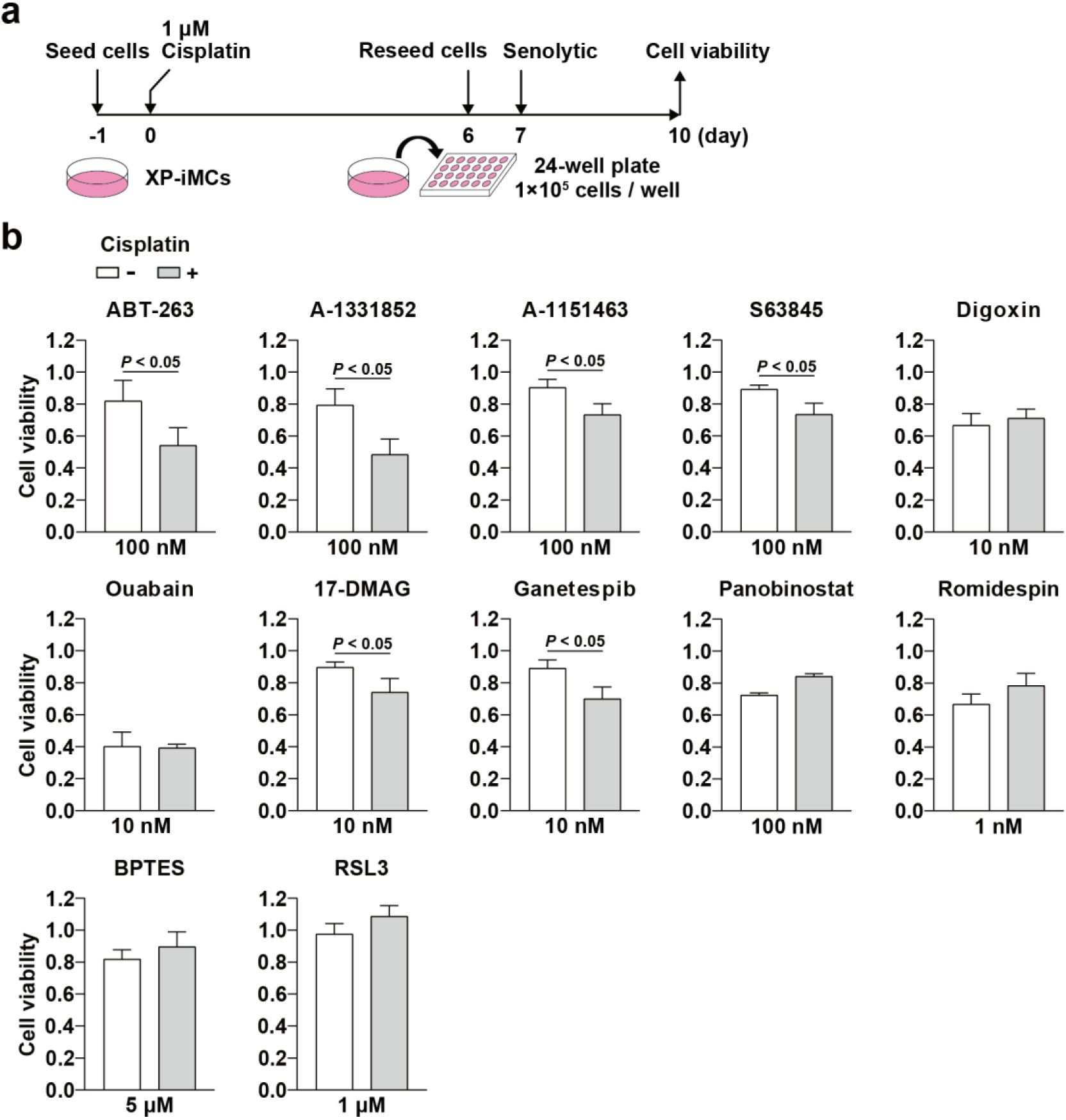
Screening of effective senolytic agents targeting cisplatin-induced senescent XP-iMCs. **(a)** Schematic representation of the senolytic screening strategy. XP-iMCs were first treated with 1 μM cisplatin for 6 days to induce senescence, followed by a 3-day treatment with candidate senolytic agents. Cell viability was measured afterward. **(b)** Cell viability of XP-iMCs treated or untreated with cisplatin, assessed after 3 days of culture with the indicated senolytic agents. Data represent means and standard deviations (n=3). Statistical significance was evaluated using a two-sided Student’s *t*-test.

### JAK inhibitors and curcuminoids as effective senomorphic agents

We sought to identify effective senomorphic agents that inhibit the accumulation of senescent XP-iMCs treated with cisplatin. For this purpose, XP-iMCs were simultaneously treated with cisplatin and senomorphic agents, as depicted in the schematic procedure (**Figure 5a**). We predicted that the simultaneous treatment would suppress SASP factor production, thereby inhibiting paracrine senescence. Fifteen senomorphic agents were chosen based on previous reports of their effects on senescent cells and their mechanisms of action (Di Micco *et al*., 2021; Wang *et al*., 2024a) (**Supplementary Table 2**). In the screening procedure, SPiDER-βGal staining combined with flow cytometric analysis was again employed, allowing for a more objective estimation of the senescent cell population (**Figure 5b**; **Supplementary Figure 3**).

**Figure 5.**
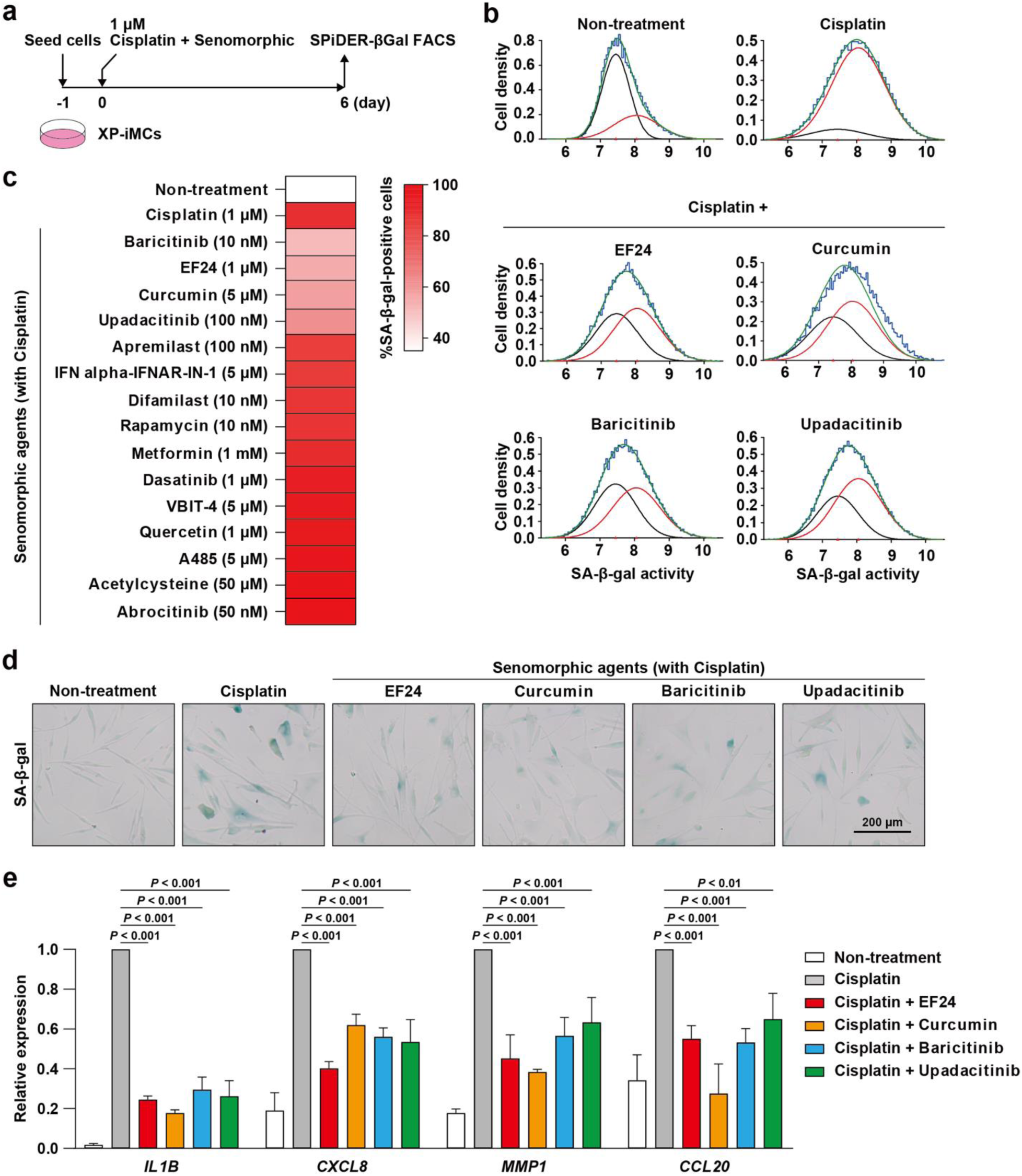
Screening for effective senomorphic agents against cisplatin-induced senescent XP-iMCs. **(a)** Schematic representation of the senomorphic screening procedure. XP-iMCs were simultaneously treated with cisplatin and senomorphic agents for 6 days, followed by SPiDER-βGal staining and FACS analysis. **(b)** Reduction of cisplatin-induced senescent cell populations by senomorphic treatment. The percentages of the two populations, SA-β-gal-positive senescent and negative non-senescent cells, were estimated as described in **Supplementary Figure 3**. **(c)** Identification of effective senomorphic agents against cisplatin-treated XP-iMCs. The percentages of SA-β-gal-positive senescent cells after the treatment with the indicated senomorphic agents were estimated as described in **panel b**. The additional results from different agent concentrations are shown in **Supplementary Figure 5**. **(d)** Effect of senomorphic agents on cisplatin-treated XP-iMCs. XP-iMCs were treated with cisplatin or simultaneously with cisplatin and the indicated senomorphic agents, followed by SA-β-gal staining. Untreated cells serve as a control. **(e)** Down-regulation of SASP genes in XP-iMCs treated with cisplatin and senomorphic agents. RT-qPCR was performed to assess the expression of the indicated SASP genes. Expression levels in XP-iMCs treated with cisplatin alone were set to 1. Data represent means and standard deviations (n=3). Statistical significance was evaluated using one-way ANOVA and Tukey’s test.

We first assessed the senescent cell population treated with cisplatin and senomorphic agents at three different concentrations (**Supplementary Figure 5**). The optimal concentrations, which yielded the lowest senescent cell populations, were selected for further experiments. These experiments were repeated to ensure reproducibility. One set of the biological replicates is shown as an example (**Figure 5c**). We observed that simultaneous treatment with cisplatin and the senomorphic agents, baricitinib, EF24, curcumin, and upadacitinib, reduced the senescent cell populations of XP-iMCs compared to cisplatin treatment alone (**Figure 5b-d**). Since baricitinib and upadacitinib are known as JAK inhibitors (Chovatiya and Paller, 2021), and EF24 and curcumin are classified as curcuminoids (He et al., 2018; Nelson et al., 2017), our results suggest that JAK inhibitors and curcuminoids are effective senomorphic agents against cisplatin-treated XP-iMCs. Since these four senomorphic agents reduced senescent cell populations of XP-iMCs treated with cisplatin, we further examined SASP gene expression after the same treatment. We found that the expression of SASP genes, which were up-regulated by cisplatin, was significantly decreased by dual treatment with cisplatin and the senomorphic agents (**Figure 5e**). These results demonstrate that the JAK inhibitors and curcuminoids can effectively inhibit the accumulation of senescent XP-iMCs induced by cisplatin treatment, thus holding potential as therapeutic agents for XP.

### Suppression of senescence-related pathways by senomorphic treatment

To understand how treatment with cisplatin and senomorphic agents affects gene expression in XP-iMCs, we performed RNA-seq as shown in **Figure 6a**. The biological replicates of RNA-seq samples clustered closely in the principal component analysis (PCA) plot, indicating a high reproducibility of the RNA-seq experiments (**Figure 6b**). Based on the results in **Figure 5**, we hypothesized that cisplatin treatment activates senescence-related pathways, while certain senomorphic agents (EF24, curcumin, baricitinib, and upadacitinib) suppress them. To test this hypothesis, we identified the differentially expressed genes (DEGs) by comparing gene expression profiles of XP-iMCs treated with cisplatin alone or in combination with senomorphic agents to non-treatment controls (**Figure 6c**; **Supplementary Figure 6**). Among the DEGs, we identified 314 genes that were commonly up-regulated and 164 genes that were commonly down-regulated across all drug-treated conditions (**Supplementary Figure 7a**). Pathway enrichment analysis revealed that genes up-regulated by cisplatin treatment were enriched in several senescence-related pathways, including p53 signaling, cytokine-cytokine receptor interaction, and PI3K/AKT signaling (**Supplementary Figure 7b**). In contrast, genes commonly up-regulated across all conditions were enriched in fewer senescence-related pathways (**Supplementary Figure 7b**). Additionally, more SASP genes were included among cisplatin-upregulated genes compared to the commonly up-regulated genes (**Supplementary Figure 7b**). These results suggest that cisplatin activates senescence-related genes and pathways, while the senomorphic agents partially suppress this activation. Conversely, genes down-regulated by cisplatin were primarily associated with lipid metabolism, alterations in which are a known feature of senescent cells (Zeng et al., 2024) (**Supplementary Figure 7c**). The commonly down-regulated genes were less enriched in these pathways, further supporting the idea that the senomorphic agents suppress the effects induced by cisplatin.

**Figure 6.**
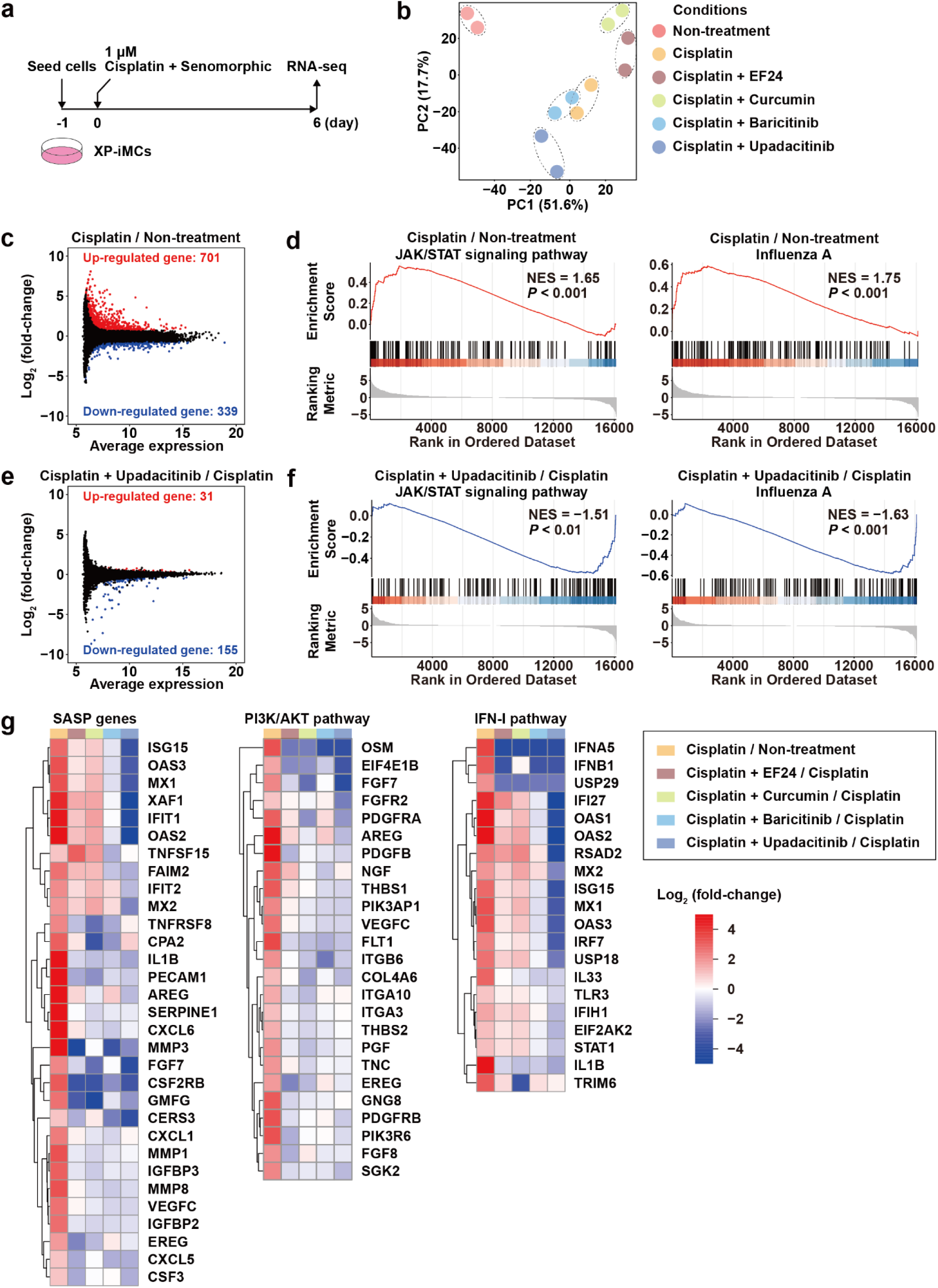
Down-regulation of senescence-related pathways by senomorphic treatment. **(a)** Schematic representation of the cisplatin and senomorphic treatment followed by RNA-seq. **(b)** PCA plot of RNA-seq data. Two biological replicates per condition were sequenced and plotted. Samples from the same condition are circled together. **(c)** The log_2_ (fold-change) in gene expression between cisplatin treatment and the non-treatment control was plotted against the average expression of each gene. Differentially expressed genes (DEGs) were defined based on a False Discovery Rate (FDR) < 0.05 and log_2_ (fold-change) > 0.7 or < –0.7. Significantly up-and down-regulated genes are represented as red and blue dots, respectively. **(d)** GSEA of the JAK/STAT signaling and Influenza A pathways, comparing expression profiles from XP-iMCs with and without cisplatin treatment. The normalized Enrichment Score (NES) quantifies the enrichment of a gene set within a ranked gene list, normalized for gene set size. **(e)** The log_2_ (fold-change) in gene expression between cisplatin + upadacitinib dual treatment and cisplatin treatment alone was plotted against the average expression of each gene. DEGs were defined based on an FDR < 0.05 and log_2_ (fold-change) > 0.3 or < –0.3. **(f)** GSEA for the JAK/STAT signaling and Influenza A pathways, comparing expression profiles from XP-iMCs treated with cisplatin + upadacitinib to those treated with cisplatin alone. **(g)** Heatmaps depicting expression alterations of SASP genes (left), PI3K/AKT pathway genes (middle), and IFN-I pathway genes (right) between the indicated conditions. Genes are clustered based on expression similarities, with red and blue indicating increased and reduced expression, respectively, between the five paired conditions compared.

### JAK/STAT, IFN-I, and PI3K/AKT pathways targeted by senomorphic agents

To understand how senomorphic agents modulate senescence pathways in cisplatin-treated XP-iMCs, we performed GSEA and observed that genes in the JAK/STAT signaling and Influenza A pathways tended to be up-regulated by cisplatin treatment (**Figure 6d**; **Supplementary Figure 7d**). This result aligns with the observation that dual treatment with cisplatin and the JAK inhibitor (baricitinib, and upadacitinib) reduces the senescent cell population compared to cisplatin treatment alone (**Figure 5**). Many of the pathways significantly enriched with cisplatin-activated genes also contained genes from the IFN-I pathway (**Supplementary Figure 7b; Supplementary Table 3**). For instance, the top-ranked Influenza A pathway consists of 33 genes, 10 of which belong to the IFN-I pathway. In this regard, the IFN-I pathway is known to be activated in senescent cells and plays a key role in regulating SASP gene expression (De Cecco et al., 2019; Wang *et al*., 2024b). Moreover, JAK/STAT signaling enhances the transcription of IFN-I response genes (Mazewski et al., 2020). Therefore, our results, in conjunction with previous studies, suggest that the JAK/STAT and IFN-I pathways are key mediators of senescence in cisplatin-treated XP-iMCs.

We next compared gene expression profiles between XP-iMCs treated with cisplatin + senomorphic agents and those treated with cisplatin alone (**Figure 6e**; **Supplementary Figure 8a**). Our analysis revealed shared up-and down-regulated genes between curcumin and EF24 treatment (**Supplementary Figure 8b**). However, no significantly enriched pathways were identified from the common DEGs. In contrast, genes up-regulated by cisplatin + upadacitinib treatment compared to cisplatin treatment were enriched in the fatty acid metabolism pathway, suggesting that lipid metabolism, which is suppressed by cisplatin treatment (**Supplementary Figure 7c**), is partly restored by upadacitinib treatment (**Supplementary Figure 8c**). Moreover, genes down-regulated by cisplatin + upadacitinib treatment were enriched in several pathways containing IFN-I pathway genes (**Supplementary Figure 8d; Supplementary Table 3**).

GSEA revealed that genes in the JAK/STAT signaling and Influenza A pathways, which are frequently up-regulated by cisplatin treatment, tended to be down-regulated by cisplatin + upadacitinib treatment (**Figure 6d, f**). Notably, the genes down-regulated by upadacitinib were significantly enriched in pathways containing the IFN-I pathway genes (**Supplementary Figure 8e**). Furthermore, many SASP genes activated by cisplatin were often down-regulated by cisplatin + senomorphic dual treatment, although SASP gene response varied between the JAK inhibitor (baricitinib, and upadacitinib) and curcuminoids (EF24 and curcumin) (**Figure 6g**). To identify key pathways targeted by the senomorphic agents, we examined expression changes of genes in pathways enriched with cisplatin-activated genes (**Supplementary Figure 7b**). Among these pathways, many genes in the PI3K/AKT signaling pathway were generally down-regulated by all four senomorphic agents, suggesting that the PI3K/AKT signaling pathway might be a common target (**Figure 6g**). In the IFN-I pathway, four genes (*IFNA5*, *IFNB1*, *USP29*, and *IL1B*) were down-regulated by the four senomorphic agents. However, other genes exhibited differential responses: many IFN-I pathway genes were up-regulated by the curcuminoids, whereas they were down-regulated by the JAK inhibitors, especially by upadacitinib (**Figure 6g**). Overall, these results suggest that the JAK inhibitors have broader effects, targeting SASP genes, PI3K/AKT signaling pathway genes, and IFN-I pathway genes, while the curcuminoids modulate a subset of genes within the same pathways.

### UV and cisplatin treatment up-regulates shared senescence-related genes

Our goal in this study was to determine whether the senomorphic agents identified through cisplatin-based screening are also effective on UV-irradiated XP-iMCs. Before addressing this question, we conducted gene expression profiling analysis of XP-iMCs, with and without UV irradiation, using HC-iMCs as a control (**Figure 7a, b**). Our hypothesis was that if the same genes were activated or repressed by cisplatin treatment and UV irradiation, we could expect similar effects from the senomorphic agents. Notably, UV irradiation altered the expression of more than 7,000 genes in XP-iMCs, whereas it affected significantly fewer genes in HC-iMCs, indicating that HC-iMCs are more resistant to UV irradiation than XP-iMCs (**Figure 7c; Supplementary Figure 9**). Additionally, expression alterations were observed between XP-iMCs and HC-iMCs even in the absence of UV irradiation. While the underlying cause remains unclear, we speculate that XP-iMCs may be more susceptible to environmental factors, such as temperature and oxygen concentration during culture.

**Figure 7.**
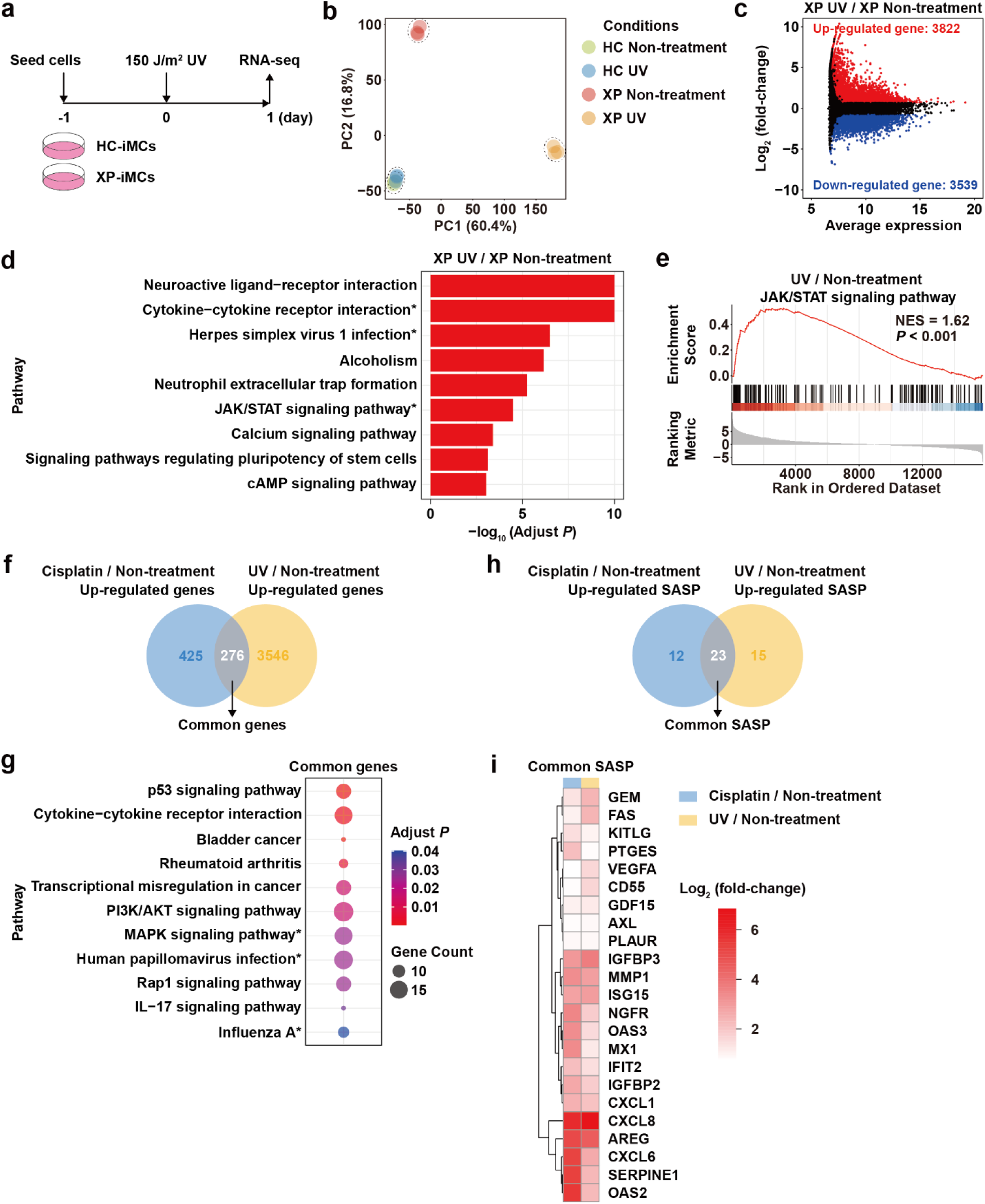
IFN-I pathway genes commonly affected by cisplatin treatment and UV irradiation. **(a)** Schematic representation of UV treatment followed by RNA-seq. **(b)** PCA plot of RNA-seq data. Three biological replicates per condition were plotted, with samples from the same condition circled together. **(c)** The log_2_ (fold-change) in gene expression between XP-iMCs with and without UV irradiation was plotted against the average expression of each gene. DEGs were defined based on an FDR < 0.05 and log_2_ (fold-change) > 0.7 or < –0.7. Significantly up- and down-regulated genes are plotted as red and blue dots, respectively. **(d)** GSEA of the indicated pathways, comparing expression profiles from XP-iMCs with and without UV irradiation. Red bars represent pathways significantly enriched with up-regulated genes in XP-iMCs with UV irradiation compared to the non-treatment control. **(e)** GSEA of the JAK/STAT signaling pathway, comparing expression profiles from XP-iMCs with and without UV irradiation. **(f)** Venn diagram showing the overlap of genes up-regulated by cisplatin treatment and UV irradiation. The overlapped genes (n=276) are referred to as common genes up-regulated by both treatments. **(g)** KEGG pathway enrichment analysis of the common genes identified in **panel f**. **(h)** Venn diagram showing the overlap of SASP genes up-regulated by cisplatin treatment and UV irradiation. The overlapped SASP genes (n=23) are referred to as common SASP genes up-regulated by both treatments. **(i)** Heatmap showing expression alterations of the common SASP genes identified in **panel h**, comparing cisplatin treatment and UV irradiation to non-treatment controls. In **panels d** and **g**, asterisks indicate pathways containing IFN-I pathway genes. Full gene lists for the respective pathways and the IFN-I pathway genes are provided in **Supplementary Table 3**.

GSEA revealed that genes in senescence-related pathways, including cytokine-cytokine receptor interaction and JAK/STAT signaling pathways, were activated in UV-irradiated XP-iMCs (**Figure 7d, e**). Genes in these two pathways were also frequently up-regulated by cisplatin treatment, indicating that UV and cisplatin tend to up-regulate genes in similar senescence-related pathways (**Figure 6d; Supplementary Figure 7d**). To further compare gene expression profiles between UV-irradiated and cisplatin-treated XP-iMCs, we identified genes commonly up-regulated by both treatment (**Figure 7f**). Pathway enrichment analysis showed that those common genes often participate in several senescence-related pathways, including p53 signaling, cytokine-cytokine receptor interaction, PI3K/AKT signaling, MAPK signaling, and Influenza A pathways (**Figure 7g**). More than half of SASP genes were up-regulated by both cisplatin and UV (**Figure 7h, i**). Collectively, these results demonstrate that UV irradiation and cisplatin treatment activate shared target genes in senescence-related pathways.

### JAK inhibitors and curcuminoids as effective senomorphic agents against UV-irradiated XP-iMCs

Since UV irradiation and cisplatin treatment affect shared target genes, we next investigated whether the senomorphic agents identified in cisplatin-based screening are also effective on UV-irradiated XP-iMCs. As our initial UV irradiation condition (150 J/m²) induced fewer than 60% SA-β-gal-positive cells (**Figure 1d**), we optimized the UV dose. To this end, we examined the effects of different UV doses on XP-iMC morphology and colony formation (**Supplementary Figure 10a, b**). We found that XP-iMCs exposed to 30 or 40 J/m^2^ UV survived at high rates and exhibited branched cell morphology by day 6, indicative of senescent melanocytes (Victorelli *et al*., 2019). Moreover, SA-β-gal-positive senescent cells were frequently detected on days 3 and 6 after 30 or 40 J/m^2^ UV irradiation (**Supplementary Figure 10c, d**). Consistently, SASP genes were up-regulated under these conditions (**Supplementary Figure 10e**). Among the conditions tested, 40 J/m^2^ UV irradiation resulted in the highest proportion of SA-β-gal-positive senescent cells.

Finally, to assess the effectiveness of senomorphic agents against UV-treated XP-iMCs, we combined a relatively low dose of UV irradiation (40 J/m^2^) with senomorphic treatment (**Figure 8a**). After three and six days of treatment with senomorphic agents, the percentages of SA-β-gal-positive cells were significantly reduced compared to untreated XP-iMCs (**Figure 8b, c**). These results demonstrate that the senomorphic agents identified through cisplatin-based screening can also suppress UV-induced senescence. Furthermore, we found that IFN-I pathway genes were significantly down-regulated by treatment with the JAK inhibitors baricitinib and upadacitinib (**Figure 8d, e**). Since IFN-I pathway genes tend to be activated by cisplatin treatment and UV irradiation and repressed by the JAK inhibitors (**Figures 6g, 8d, e**), our results suggest that the JAK inhibitors inhibit UV-induced senescence of XP-iMCs through suppression of the IFN-I pathway (see Discussion).

**Figure 8.**
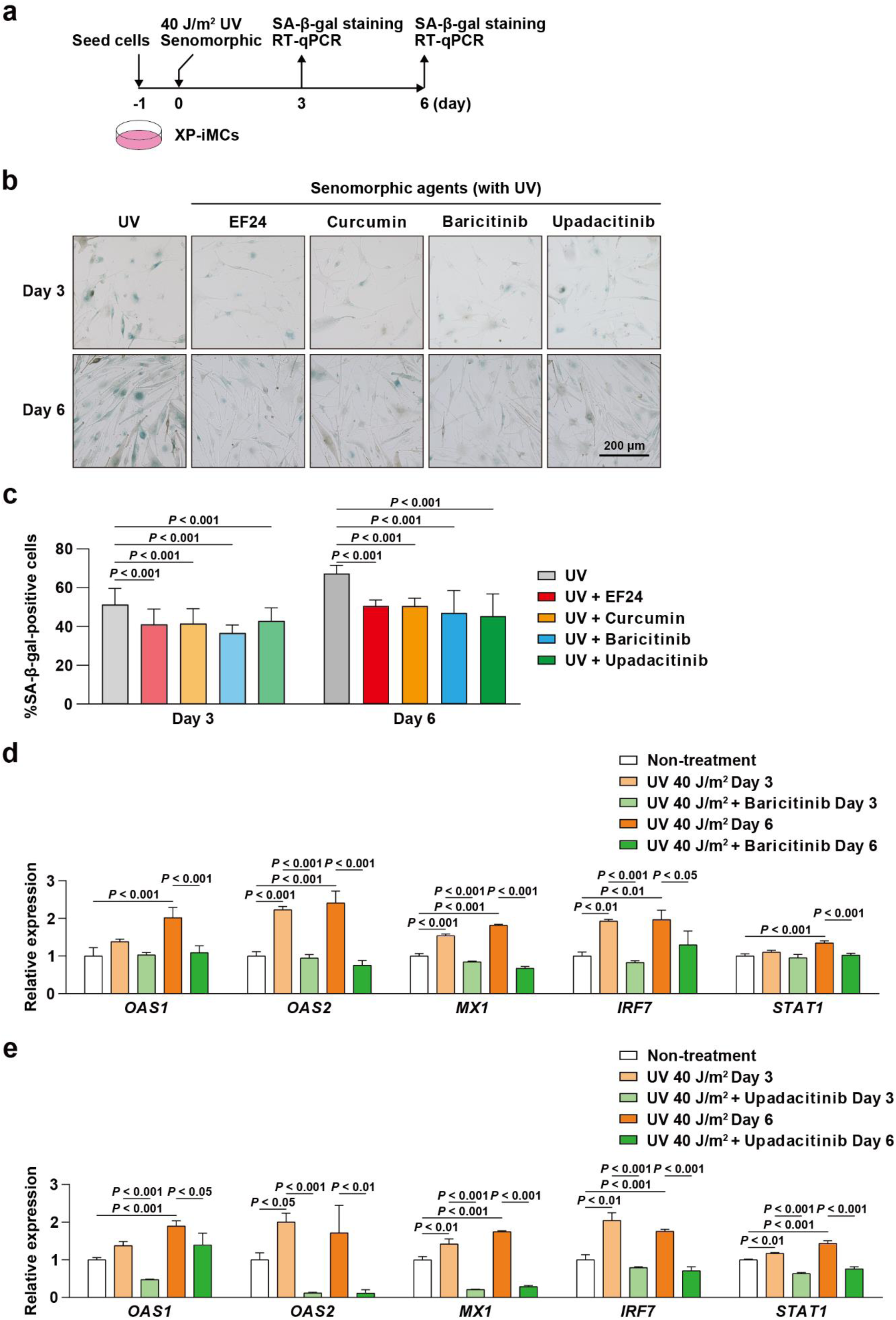
Effects of senomorphic agents on UV-irradiated XP-iMCs. **(a)** Schematic representation of the UV irradiation procedure and experimental design. **(b)** SA-β-gal staining of UV-irradiated XP-iMCs treated with the indicated senomorphic agents. UV-irradiated XP-iMCs without senomorphic treatment are shown as a control. **(c)** Quantification of SA-β-gal-positive XP-iMCs irradiated with UV, with and without senomorphic treatment. SA-β-gal staining images, as exemplified in **panel b**, were quantified. **(d)** Down-regulation of IFN-I pathway genes in UV-irradiated XP-iMCs treated with baricitinib. **(e)** Down-regulation of IFN-I pathway genes in UV-irradiated XP-iMCs treated with upadacitinib. In **panels c-e**, data represent means and standard deviations are shown (n=3). Statistical significance was assessed using one-way ANOVA and Tukey’s test.

## Discussion

In this study, we demonstrated that UV irradiation induces senescence of XP-iMCs, which are melanocytes derived from an XP patient. Additionally, we developed a cisplatin-based drug screening system to identify effective senotherapeutic agents for the treatment of XP. Cisplatin was selected to mimic UV-induced senescence due to its high reproducibility and easily adjustable dosage. Through this screening, we first identified two groups of effective senolytic agents against cisplatin-treated XP-iMCs: BCL-2-like protein inhibitors (ABT-263, A-1331852, A-1155463, and S63845) and HSP90 inhibitors (17-DMAG and ganetespib). However, we also observed that melanocytes are inherently more vulnerable to senolytic agents than keratinocytes, even in the absence of cisplatin treatment. This finding led us to prioritize senomorphic agents as a potentially safer option for the treatment of XP, although optimizing the dosage of senolytic agents may still hold therapeutic potential.

Senomorphic agents have recently been shown to extend lifespan and reduce age-related inflammation (Di Micco *et al*., 2021; Xu *et al*., 2015). Among the senomorphic agents we tested, JAK inhibitors, baricitinib and upadacitinib, effectively reduced the percentages of senescent XP-iMCs and SASP genes expression. Expression profiling further revealed that the JAK/STAT pathway is activated by both UV irradiation and cisplatin treatment. This finding aligns with a previous report indicating that activation of the JAK/STAT pathway is a common hallmark of age-related diseases (Liu *et al*., 2019). Moreover, coactivation of the JAK/STAT, PI3K/AKT, and IFN-I pathways has been observed under various conditions, including oxidative stress, antiviral response, and carcinogenesis (Lu et al., 2008; Mazewski *et al*., 2020). Our expression analysis also indicates that genes in the PI3K/AKT and IFN-I pathways are down-regulated by treatment with the JAK inhibitors. Based on these results, we propose that the activation of the JAK/STAT, PI3K/AKT, and IFN-I pathways plays a critical role in the etiology of XP. Notably, the JAK inhibitors can simultaneously suppress all these pathways, highlighting their potential as therapeutic agents for XP.

Additionally, curcuminoids (curcumin and its analog EF24) were also found to suppress senescence-related pathways in the drug screening. While the mechanisms of curcuminoids are complex, previous studies have shown that curcuminoids can inhibit NF-κB signaling, which functions downstream of the PI3K/AKT pathway (Marquardt et al., 2015; Yin *et al*., 2016). Our study suggests that the PI3K/AKT pathway is a common target for both the JAK inhibitors and curcuminoids. As a natural product, curcumin might be a safer option for XP treatment. On the other hand, several JAK inhibitors have already been approved by the FDA for treating certain diseases, making them promising candidates for repurposing in XP treatment (Chovatiya and Paller, 2021; Liu *et al*., 2019; Xu *et al*., 2015). Furthermore, since JAK inhibitors are available in topical formulations, they could be applied in low doses to healthy individuals to mitigate UV-induced skin damage after sunburn.

In this study, we employed cisplatin as a senescence inducer to mimic UV irradiation. Our findings indicate that both cisplatin treatment and UV irradiation activate similar senescence-related pathways, including the JAK/STAT and PI3K/AKT pathways, while up-regulating multiple shared SASP genes. Moreover, agents identified through cisplatin-based drug screening significantly suppressed senescence of UV-irradiated melanocytes. These results suggest that cisplatin treatment successfully recapitulates the effects of UV irradiation in our screening system. Therefore, this cisplatin-based approach holds promise for a large-scale drug screening to identify potential senotherapeutic agents for XP treatment. Finally, further investigation into therapeutics for other XP subtypes is also warranted.

## Materials and Methods

### Cell lines and culture conditions

Induced pluripotent stem cells (iPSCs) from a healthy control (HC-iPSCs) and a xeroderma pigmentosum patient carrying a single nucleotide variation in the *XPA* gene (XP-iPSCs) were generated using TIG-120 (normal diploid fibroblasts from human skin; JCRB0542) and XP3OS cells (fibroblasts from xeroderma pigmentosum patient; JCRB0303), respectively, as previously described (Hosaka *et al*., 2019; Kondo and Yonezawa, 1995; Takebe et al., 1977; Takemori *et al*., 2024). Both cell lines were obtained from the Japanese Collection of Research Bioresources Cell Bank. XP3OS cells harbor a Japanese founder single nucleotide variation in the *XPA* gene (Nishigori et al., 1994; Satokata *et al*., 1990; Takebe *et al*., 1977).

The established melanocyte lines, HC-iMCs and XP-iMCs, were cultured in M254 medium (Thermo Fisher Scientific, M254500) supplemented with human melanocyte growth supplement (Thermo Fisher Scientific, S0025) and 1% Penicillin-Streptomycin-Amphotericin B solution (Wako, 161-23181). Culture dishes were coated with fibronectin derived from human plasma (Merk, F0895). Normal human epidermal melanocytes (NHEM) were also cultured under the same condition. The hTERT-immortalized human keratinocyte line (Ker-CT) was cultured in KBM-Gold^TM^ keratinocyte basal medium (Lonza, 00192151) supplemented with KGM-Gold^TM^ keratinocyte SingleQuots^TM^ growth factors (Lonza, 00192152). Cells were cultured at 37℃ in a CO_2_/multi-gas incubator (Astec, SMA-80) under 5% CO_2_ and 3% O_2_ conditions.

### Colony formation

Colony formation assays were performed as previously described (Franken et al., 2006). Briefly, XP-iMCs were irradiated with UV and cultured for six days. Cells were washed twice with PBS to remove the media, fixed with 10% neutral buffered formalin for 15 minutes at room temperature, washed once with PBS, and stained with 0.5% crystal violet for 30 minutes at room temperature.

### Cell viability

Cells were seeded in 96-well cell culture plates at a density of 10,000 cells per well in M254 medium and cultured for one day to allow the cells to adhere to the culture plates. The cells were exposed to various concentrations of cisplatin (Selleck, S1166) in M254 medium and maintained for an additional 72 hours. The cell numbers were measured using the Synergy LX Multimode Reader (Agilent Technologies) and the Cell Count Reagent SF Kit (Nacalai Tesque, 07553-44). Cisplatin IC_50_ values were estimated from three independent experiments using a nonlinear regression model.

For senolytic drug screening with sequential treatment, as depicted in **Figure 4a**, XP-iMCs were cultured in 10 cm dishes with M254 medium containing 1 μM cisplatin for six days. Afterward, the cells were reseeded in 24-well cell culture plates at a density of 100,000 cells per well and cultured for one day to allow attachment. Following this, the cells were treated with one of the senolytic agents for 72 hours before cell viability was assessed using the Cell Count Reagent SF Kit. Simultaneous treatment with cisplatin and senotherapeutic agents, including senolytic and senomorphic compounds, was performed, as shown in **Figure 5a** and **Supplementary Figure 4a**. XP-iMCs were exposed to 1 μM cisplatin and a senolytic agent at the indicated concentrations for six days, followed by cell viability measurement.

### Senotherapeutic drug screening agents

The followed agents were applied for senolytic drug screening: ABT-263 (Selleck, S1001), A-1331852 (Selleck, S7801), A-1155463 (Selleck, S7800), S63845 (MedchemExpress, HY-100741), BPTES (Selleck, S7753), RSL3 (Selleck, S8155), digoxin (Sigma, D6003), ouabain (Sigma, O3125), 17-DMAG (Selleck, S1142), ganetespib (Selleck, S1159), panobinostat (Selleck, S1030), and romidepsin (MedchemExpress, HY-15149).

The followed agents were applied for senomorphic drug screening: A485 (Selleck, S8740), acetylcysteine (Selleck, S1623), curcumin (Selleck, S1848), EF24 (MedchemExpress, HY-119272), metformin (Selleck, S5958), apremilast (Selleck, S8034), difamilast (MedchemExpress, HY-109085), abrocitinib (Selleck, S8765), baricitinib (Selleck, S2851), upadacitinib (Selleck, S8162), IFN alpha-IFNAR-IN-1 (Selleck, E2982), VBIT-4 (Selleck, S3544), rapamycin (Selleck, S1039), dasatinib (Selleck, S1021), and quercetin (Selleck, S2391).

### Western blotting

Western blotting was performed as previously described (Wang *et al*., 2024b). The primary antibodies used in this study were 1:1,000-diluted mouse anti-XPA 12F5 (Santa Cruz Biotechnology, sc-56813), 1:1,000-diluted mouse anti-Flag M2 (Sigma, F1804), 1:1,000-diluted rabbit anti-IL-6 (abcam, ab233706), 1:1,000-diluted rabbit anti-MMP1 (proteintech, 10371-2-AP), 1:1,000-diluted rabbit anti-MMP3 (Cell Signaling, #14351), 1:1,000-diluted rabbit anti-α/β-tubulin (Cell Signaling, #2148), and 1:2,000-diluted mouse anti-β-actin (Wako, 017-24551). The secondary antibodies were 1:2,500-diluted HRP-conjugated anti-mouse IgG (Promega, W4021) and 1:1,000-diluted HRP-conjugated anti-rabbit IgG (Cytiva, NA9340). Signal detection was performed using the ECL Prime Western Blotting System (Cytiva) and ImageQuant LAS 4000 mini (GE Healthcare).

### Immunofluorescence

Immunofluorescence was performed as previously described (Wang *et al*., 2024b). The primary antibodies used in this study were 1:500-diluted mouse anti-XPA 12F5 (Santa Cruz Biotechnology, sc-56813), 1:500-diluted mouse anti-Flag M2 (Sigma, F1804), 1:500-diluted mouse anti-phospho-Histone H2A.X (Ser139) clone JBW301 (Sigma-Aldrich, 05-636), and 1:500-diluted mouse anti-p21 (187) (Santa Cruz Biotechnology, sc-817). The secondary antibody was 1:500-diluted Alexa Fluor 488-conjugated goat anti-mouse IgG H&L (Abcam, ab150113). Cells were mounted on microscope slides with Antifade Mounting Medium containing DAPI (VECTASHIELD, H-1200). Immunofluorescence images were captured using a confocal microscope (Olympus, FV1000) with a 60× lens.

### Senescence-associated (SA)-β-gal staining

SA-β-gal staining of melanocytes was performed as previously described (Dimri *et al*., 1995). Briefly, cells were seeded on cover glasses and fixed with PBS containing 2% formaldehyde and 0.2% glutaraldehyde for 5 minutes at room temperature. After two washes with PBS, the cells were stained overnight at 37℃ in a non-CO_2_ incubator (EYELA, SLI-700) using X-gal staining solution [150 nM NaCl, 2 mM MgCl_2_, 5 mM K_3_Fe(CN)_6_, 5 mM K_4_Fe(CN)_6_, 40 mM Na_2_HPO_4_ pH 6.0, and 1 mg/mL X-gal (Sigma, B9146)]. Following staining, the cells were washed with PBS and water. Images were captured using an upright microscope (ZEISS, Primostar 3) with a 10× lens.

### SA-β-gal quantification

Quantification of SA-β-gal-positive cells was performed as previously described (Wang *et al*., 2024b). Briefly, melanocytes were stained using the Cellular Senescence Detection Kit SPiDER-βGal (Dojindo, SG03) and analyzed with the BD FACSCanto II Flow Cytometry System (BD Biosciences). The percentages of SA-β-gal-positive senescent and negative non-senescent cells were estimated using the “mixdist” package in R.

### RT-qPCR

Total RNA was isolated using Trizol reagent (Thermo Fisher Scientific, 15596018) and subjected to reverse transcription using the PrimeScript™ RT Master Mix (Takara Bio, RR036A) according to the manufacturer instructions. The resulting cDNA was used as a template for quantitative PCR, performed using the KAPA SYBR FAST qPCR Kit (Kapa Biosystems, KK4600) and the 7300 Real-Time PCR System (Applied Biosystems). Expression levels of target genes were normalized to the mRNA levels of the β2-microglobulin (*B2M*) or *GAPDH* gene as internal controls. The fold change in the target gene expression was calculated using the 2^-ΔΔCT^ method (Livak and Schmittgen, 2001).

The RT-qPCR primer sequences for the respective genes were as follows: *B2M* forward (5’-GGCATTCCTGAAGCTGACA-3’) and reverse (5’-CTTCAATGTCGGATGGATGAAAC-3’); *GAPDH* forward (5’-TGTTGCCATCAATGACCCCTT-3’) and reverse (5’-CTCCACGACGTACTCAGCG-3’); *IL1B* forward (5’-AGCTCGCCAGTGAAATGATGG-3’) and reverse (5’-GTCCTGGAAGGAGCACTTCAT-3’); *IL6* forward (5’-ACATCCTCGACGGCATCTCA-3’) and reverse (5’-TCACCAGGCAAGTCTCCTCA-3’); *CXCL8* forward (5’-GTTTTTGAAGAGGGCTGAGAATTC-3’) and reverse (5’-CCCTACAACAGACCCACACAATAC-3’); *MMP1* forward (5’-AATAGTGGCCCAGTGGTTGA-3’) and reverse (5’-GGCTGCTTCATCACCTTCAG-3’); *MMP3* forward (5’-CACTCACAGACCTGACTCGGTT-3’) and reverse (5’-AAGCAGGATCACAGTTGGCTGG-3’); *CCL20* forward (5’-CTCCTGGCTGCTTTGATGTC-3’) and reverse (5’-TGCTTGCTGCTTCTGATTCG-3’); *OAS1* forward (5’-GTTGCCACTCTCTCTCCTGT-3’) and reverse (5’-CACCTTGGACACACACACAG-3’); *OAS2* forward (5’-TTCTCCAGCCCAACAAATGC-3’) and reverse (5’-AGTCTTCAGAGCTGTGCCTT-3’); *MX1* forward (5’-CGGAATCTTGACGAAGCCTG-3’) and reverse (5’-CCTTTCCTTCCTCCAGCAGA-3’); *IRF7* forward (5’-CACACACACATGCTGGACTC-3’) and reverse (5’-CCTTGGTTGGGACTGGATCT-3’); *STAT1* forward (5’-TGCCACCATCCGTTTTCATG-3’) and reverse (5’-ATATTCCCCGACTGAGCCTG-3’).

### RNA sequencing

RNA-seq was performed as previously described (Wang *et al*., 2024b). Briefly, total RNA was treated with RQ1 RNase-Free DNase (Promega, M6101) and purified by phenol/chloroform extraction followed by ethanol precipitation. Poly (A) RNA was isolated using the NEBNext Poly (A) mRNA Magnetic Isolation Module (New England Biolabs, E7490S), and cDNA fragments were prepared by the NEBNext Ultra II RNA Library Prep Kit for Illumina (New England Biolabs, E7770S) and purified with AMPure XP beads (Beckman Coulter, A63880). Adaptor-ligated DNA fragments were generated using the NEBNext Ultra II Q5 Master Mix (New England Biolabs, M0544S) and NEBNext Multiplex Oligos for Illumina (Index Primers Set 1) (New England Biolabs, E7335S). The libraries were sequenced on an Illumina NextSeq2000 platform using 51 bp × 2 paired-end reads according to the manufacturer instructions.

To ensure consistent read numbers and depth across drug treatment and UV irradiation experiment series, sequencing reads from biological triplicates and remixed to duplicates by randomly selecting and evenly distributing reads from the third replicates to the other two duplicates using SeqKit v2.0.4, after confirming high consistency between all 3 replicates by principal component analysis (Shen et al., 2024) (**Figure 7b**). To minimize biases due to variations in read numbers, sequenced data from respective samples were down-sampled to 30 million reads by random selection using SeqKit v2.0.4. Reads were aligned to the human reference genome GRCh38 using STAR aligner v2.7.10a (Dobin et al., 2013). Multi-mapped reads were discarded using SAMtools (Li et al., 2009), and gene expression levels were quantified using HTSeq v2.0.3 based on the GRCh38.p14 genome reference and GENCODE v44 gene annotation (Anders et al., 2015; Frankish et al., 2021). Differentially expressed genes were identified using DESeq2 v1.40.2 (Love et al., 2014), while Kyoto Encyclopedia of Genes and Genomes (KEGG) analysis and Gene Set Enrichment Analysis (GSEA) were conducted with clusterProfiler package v4.8.2 (Wu et al., 2021).

### Statistical analysis

Data are presented as means ± standard deviations. Statistical comparisons between two groups were performed using a two-tailed Student’s *t*-test, while comparisons among more than two groups were conducted using one-way ANOVA and Tukey’s test. All statistical analyses were performed using GraphPad Prism (version 10.1.1).

### Data availability

The RNA-seq data were deposited in the Gene Expression Omnibus (GEO) database with accession number GSE293695. All relevant data supporting the findings of this study are available from the authors upon request.

## Acknowledgments

We would like to thank Chihiro Kinugasa, Kayo Tagawa, Michiko Kanbayashi, and Makiko Fukuuchi for their generous support in the laboratories. This work was supported by the Japan Agency for Medical Research and Development (AMED) under Grant Numbers JP24ym0126809j0003 and JP24ama221535h0001, KOSÉ Cosmetology Research Foundation, Atsushi Kukita Award from the Japanese Society of Pigment Cell Research, the Naito Foundation, Takeda Science Foundation, the Nakatomi Foundation, the Ichiro Kanehara Foundation, The POLA R&M Grant for Vitiligo Research, the Grant for Joint Research Program of the Institute for Genetic Medicine, Hokkaido University (to T.F.), Japan Society for the Promotion of Science (JSPS) KAKENHI grant JP20K23376, Takeda Science Foundation, the Mitsubishi Foundation, the Photo-excitonix Project in Hokkaido University (to K.N.), Japanese Government (MEXT) Scholarship, Japan Science and Technology Agency (JST) SPRING grant JPMJSP2119, and JSPS Research Fellowship for Young Scientists (DC2) (to X.W.).

