## Supplementary Figures and Tables for "Senotherapeutic potential against xeroderma pigmentosum"

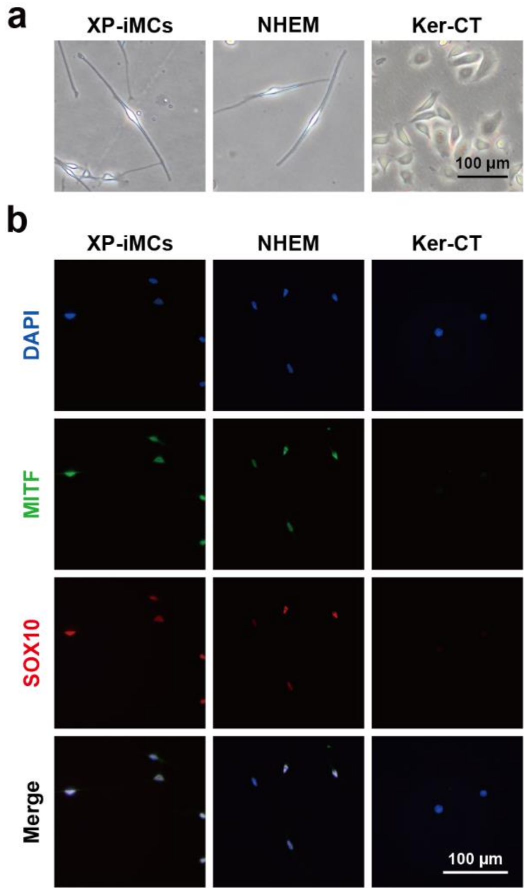


**Supplementary Figure 1. Characterization of XP-iMCs.**

**(a)** Cell morphology of XP-iMCs and NHEM melanocytes, and Ker-CT keratinocytes.

**(b)** Immunofluorescence staining of melanocyte markers MITF and SOX10 in XP-iMCs, NHEM and Ker-CT. Cell nuclei were also stained with DAPI.

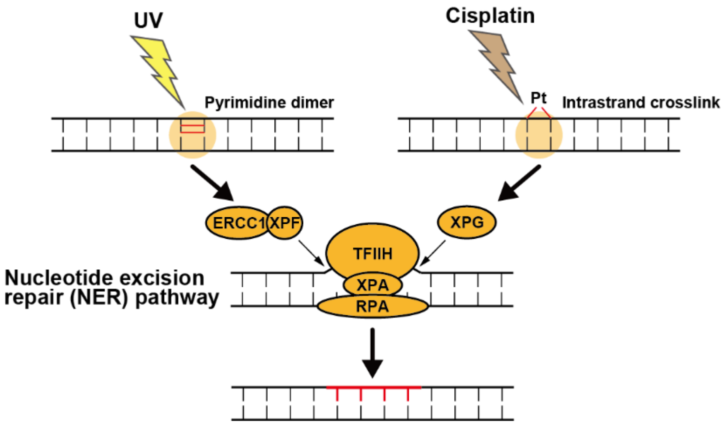


**Supplementary Figure 2. Both UV- and cisplatin-formed DNA lesions are repaired by the nucleotide excision repair (NER) pathway.**

We propose using the cisplatin-induced senescence model to identify senotherapeutic agents. This screening approach is conceived from the fact that UV-induced pyrimidine dimers and cisplatin-induced intrastrand crosslinks are repaired through the same NER pathway.

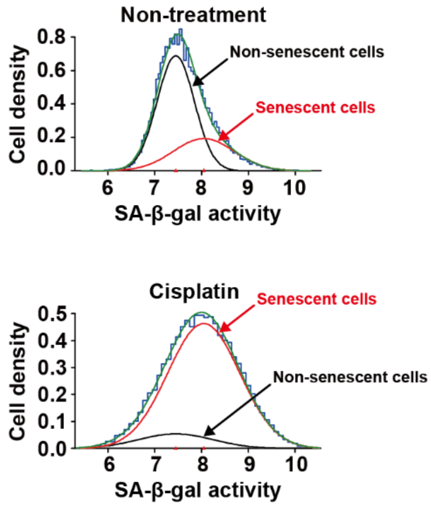


**Supplementary Figure 3. Quantification of XP-iMC senescent cells by flow cytometry.**

XP-iMCs were treated or untreated with cisplatin, and the percentage of SA-β-gal-positive cells was quantified. Flow cytometric data (blue) were used to estimate the entire cell population (green), which was in turn divided into two subset populations: SA-β-gal-positive senescent cells (red) and negative non-senescent cells (black). The percentages of these two populations were estimated using the mixdist package in R. The data shown here are representative examples corresponding to those presented in **Figure 5b**.

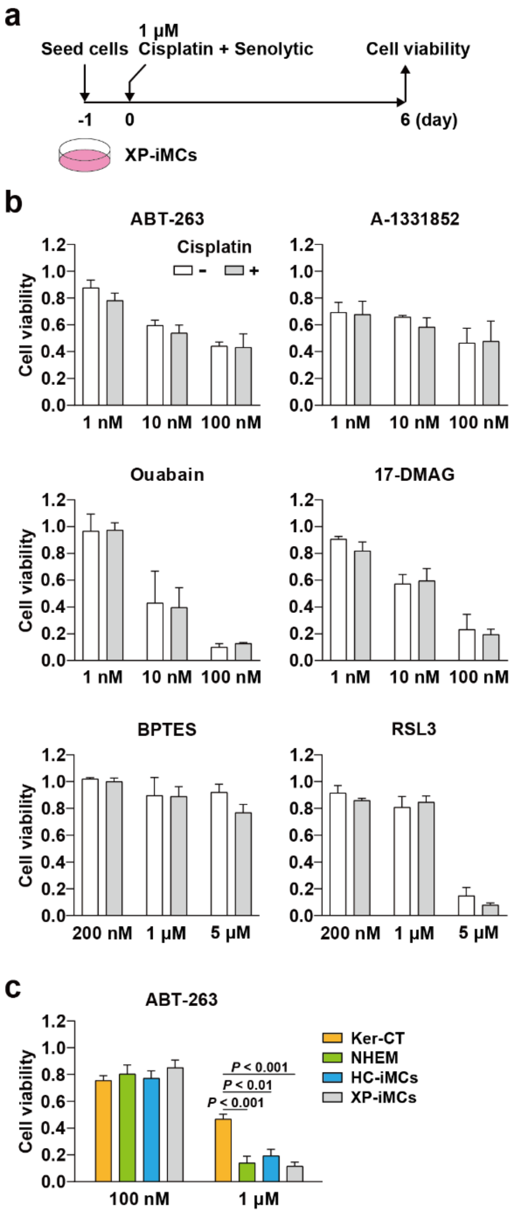


**Supplementary Figure 4. Effect of senolytic agents on XP-iMCs simultaneously treated with cisplatin.**

**(a)** Schematic representation of the senolytic screening procedure.

**(b)** Effect of senolytic agents on the viability of XP-iMCs. Cells were treated with cisplatin and senolytic agents as depicted in **panel a**, and cell viability was assessed.

**(c)** Effect of ABT-263 on the viability of keratinocytes and melanocytes. Keratinocytes (Ker-CT) and three melanocyte cell lines (NHEM, HC-iMCs, and XP-iMCs) were treated with ABT-263 at the indicated concentrations for 72 hours, followed by cell viability measurement.

In **panels b** and **c**, data represent means and standard deviations (n=3). Statistical significance was assessed by a one-way ANOVA and Tukey's test. No significant difference was observed between cisplatin treated and untreated cells in **panel b**.

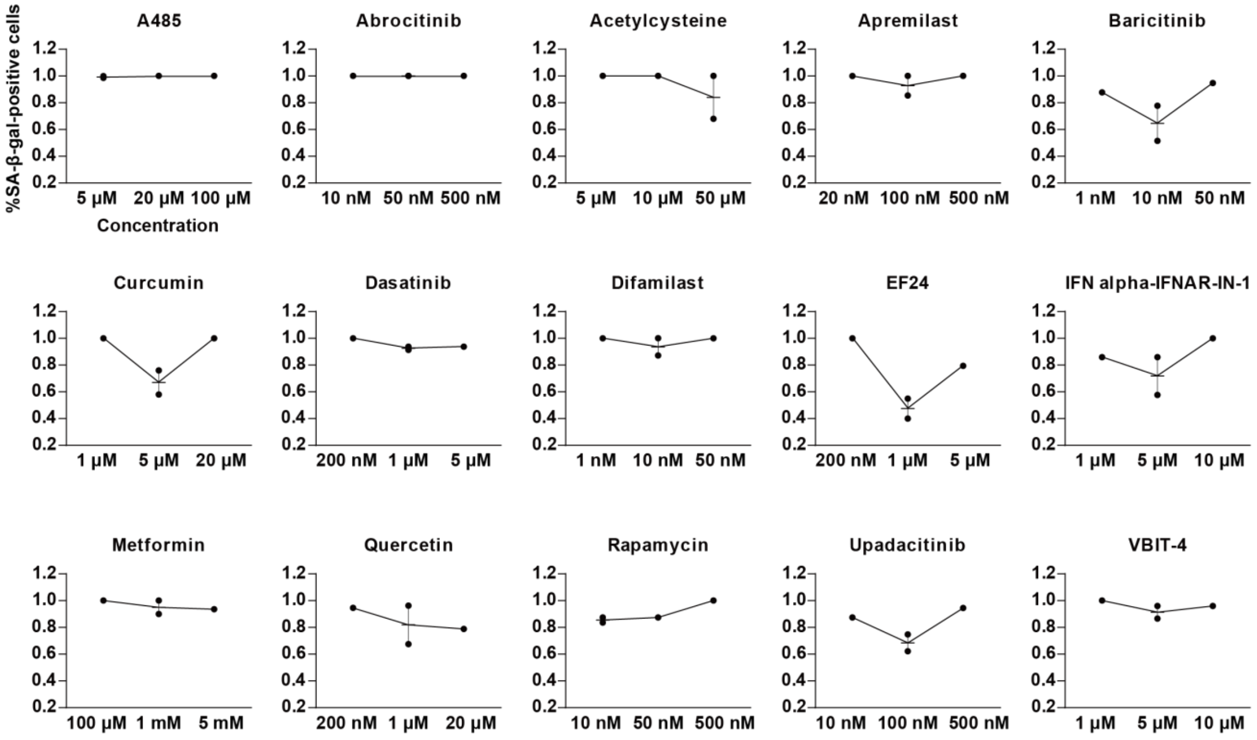


**Supplementary Figure 5. Effect of senomorphic agents on XP-iMCs treated with cisplatin.** The percentages of SA-β-gal-positive senescent XP-iMCs simultaneously treated with cisplatin and the indicated senomorphic agents were estimated as shown in **Figure 5a, b** (filled dots). In addition to the results shown in **Figure 5c**, different concentrations of the senomorphic agents were subjected to the same analysis. For respective senomorphic agents, experiments conducted at the specific concentrations that resulted in the lowest percentages of SA-β-gal-positive senescent cells were repeated to confirm reproducibility. Means are represented by short bars.

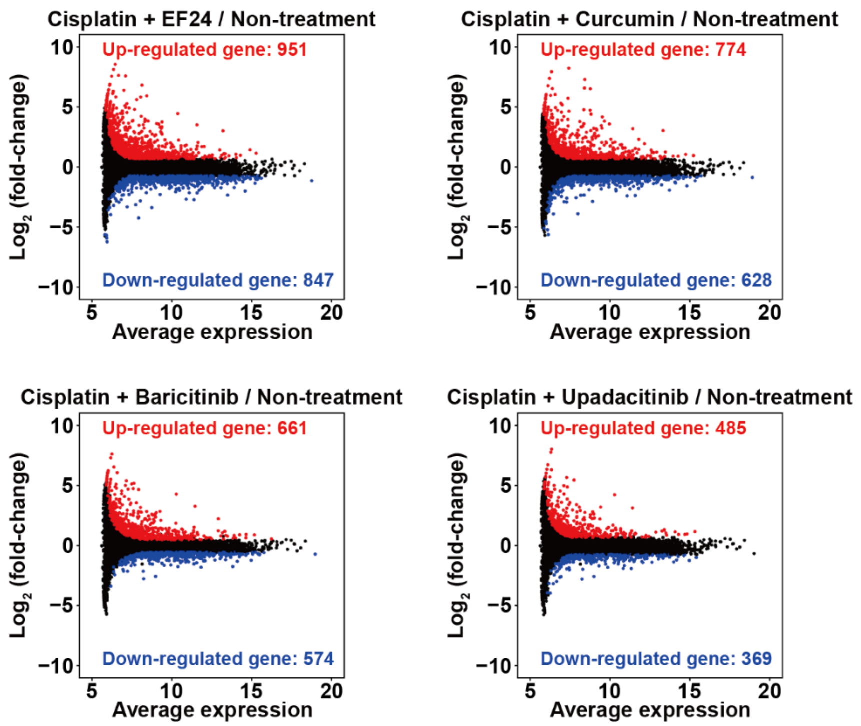


**Supplementary Figure 6. Gene expression alterations in XP-iMCs treated with cisplatin and senomorphic agents.**

The $\log_2$ (fold-change) in gene expression between cisplatin + senomorphic treatment and the non-treatment control was plotted against the average expression of each gene. Differentially expressed genes (DEGs) were identified based on an FDR < 0.05 and a $\log_2$ (fold-change) > 0.7 or < –0.7. Significantly up- and down-regulated genes are represented as red and blue dots, respectively.

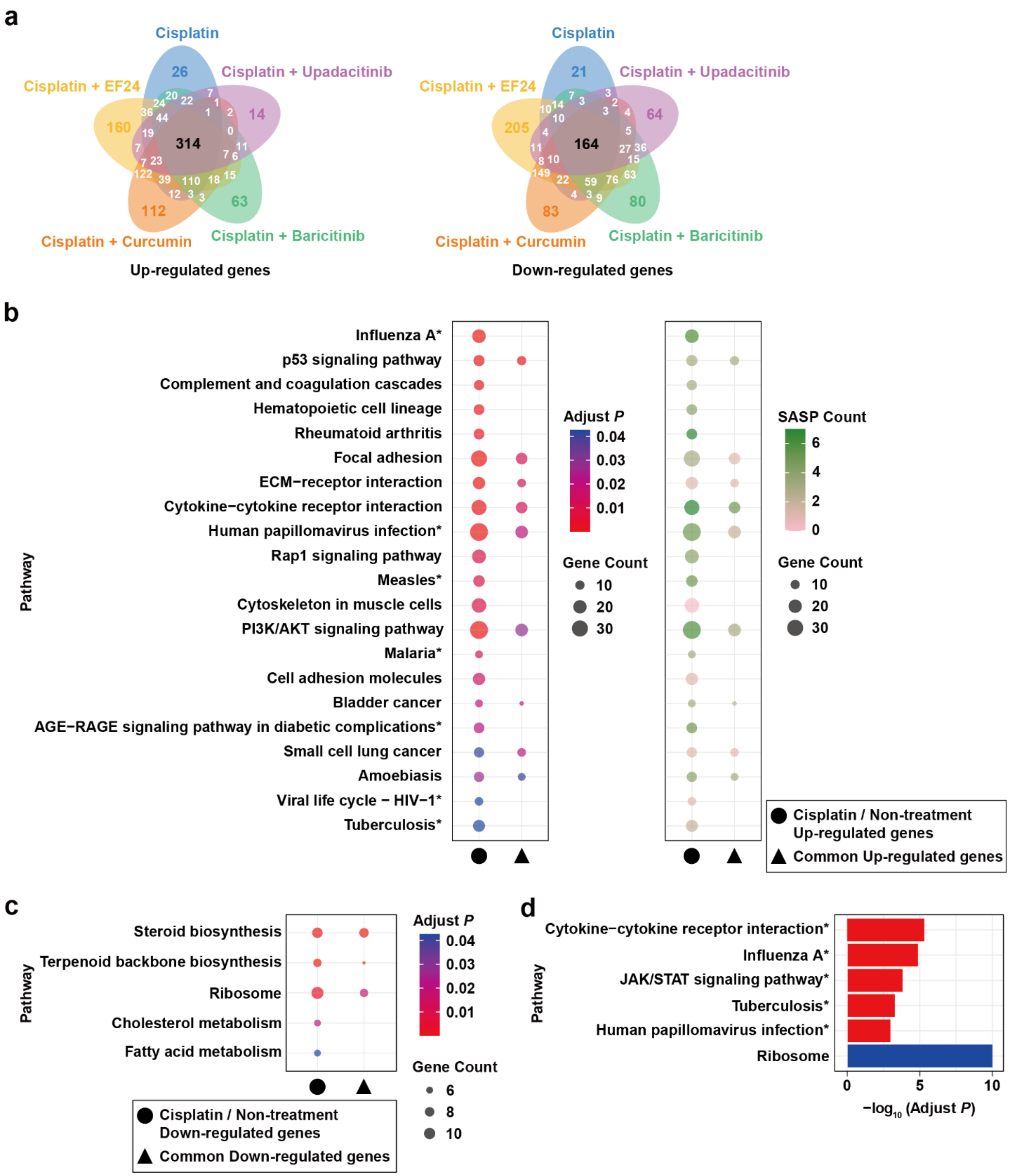


**Supplementary Figure 7. Senescence-related pathways activated in XP-iMCs treated with cisplatin.**

**(a)** Venn diagrams illustrating the overlap of significantly up- (left) or down-regulated (right) genes across the indicated treatments compared to the non-treatment control.

**(b)** KEGG pathway enrichment analyses of genes up-regulated by cisplatin treatment and the common up-regulated genes (n=314), as defined in **panel a**. The color in the left panel indicates the adjusted *p*-value for each pathway, while the color in the right panel reflects the number of SASP genes involved in each pathway.

**(c)** KEGG pathway enrichment analyses of genes down-regulated by cisplatin treatment and the common down-regulated genes (n=164), as defined in **panel a**.

**(d)** GSEA of genes in the indicated pathways. Gene expression profiles from XP-iMCs with and without cisplatin treatment were used in this analysis. Red and blue bars represent pathways significantly enriched with up- and down-regulated genes, respectively, by cisplatin treatment.

In **panels b** and **d**, asterisks indicate pathways containing IFN-I pathway genes. Full gene lists for the respective pathways and the IFN-I pathway genes are provided in **Supplementary Table 3**.

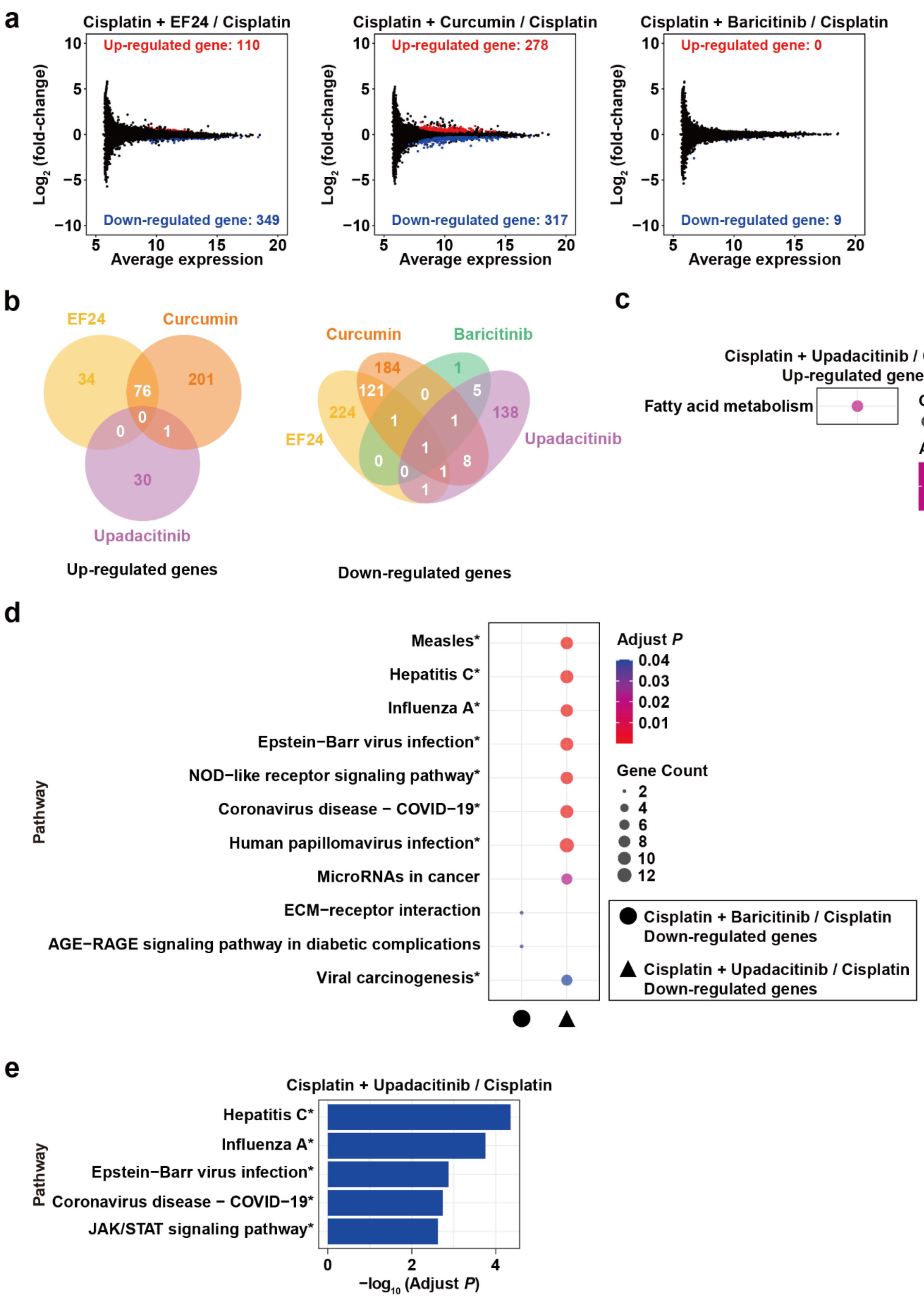
a
Cisplatin + EF24 / Cisplatin
Up-regulated gene: 110
Down-regulated gene: 349
Cisplatin + Curcumin / Cisplatin
Up-regulated gene: 278
Down-regulated gene: 317
Cisplatin + Baricitinib / Cisplatin
Up-regulated gene: 0
Down-regulated gene: 9
Log2 (fold-change)
Average expression
b
EF24
Curcumin
Upadacitinib
Up-regulated genes
Curcumin
Baricitinib
EF24
Upadacitinib
Down-regulated genes
c
Cisplatin + Upadacitinib / Cisplatin
Up-regulated genes
Fatty acid metabolism
Gene Count
Adjust P
0.042
d
Measles*
Hepatitis C*
Influenza A*
Epstein−Barr virus infection*
NOD−like receptor signaling pathway*
Coronavirus disease − COVID−19*
Human papillomavirus infection*
MicroRNAs in cancer
ECM−receptor interaction
AGE−RAGE signaling pathway in diabetic complications
Viral carcinogenesis*
Pathway
Adjust P
Gene Count
Cisplatin + Baricitinib / Cisplatin
Down-regulated genes
Cisplatin + Upadacitinib / Cisplatin
Down-regulated genes
e
Cisplatin + Upadacitinib / Cisplatin
Hepatitis C*
Influenza A*
Epstein−Barr virus infection*
Coronavirus disease − COVID−19*
JAK/STAT signaling pathway*
Pathway
−log10 (Adjust P)

**Supplementary Figure 8. Down-regulation of IFN-I pathway genes in XP-iMCs treated with the JAK inhibitors baricitinib and upadacitinib.**

**(a)** The $\log_2$ (fold-change) in gene expression between cisplatin + senomorphic treatment and cisplatin treatment alone was plotted against the average expression of each gene. DEGs were defined based on an FDR < 0.05 and $\log_2$ (fold-change) > 0.3 or < –0.3. Significantly up- and down-regulated genes are represented as red and blue dots, respectively.

**(b)** Venn diagrams illustrating the overlap of significantly up- (left) or down-regulated (right) genes across the indicated dual treatments compared to cisplatin treatment alone.

**(c)** KEGG pathway enrichment analysis of the up-regulated genes in XP-iMCs treated with upadacitinib.

**(d)** KEGG pathway enrichment analysis of the down-regulated genes in XP-iMCs treated with baricitinib or upadacitinib.

**(e)** GSEA of genes in the indicated pathways. Gene expression profiles from XP-iMCs treated with cisplatin + upadacitinib and treated with cisplatin treatment alone were used in this analysis. Blue bars represent pathways significantly enriched with down-regulated genes in XP-iMCs with dual treatment (cisplatin + upadacitinib) when comparing to cisplatin treatment alone.

In **panels d** and **e**, asterisks indicate pathways containing IFN-I-related genes. Full gene lists for the respective pathways and the IFN-I pathway genes are provided in **Supplementary Table 3**.

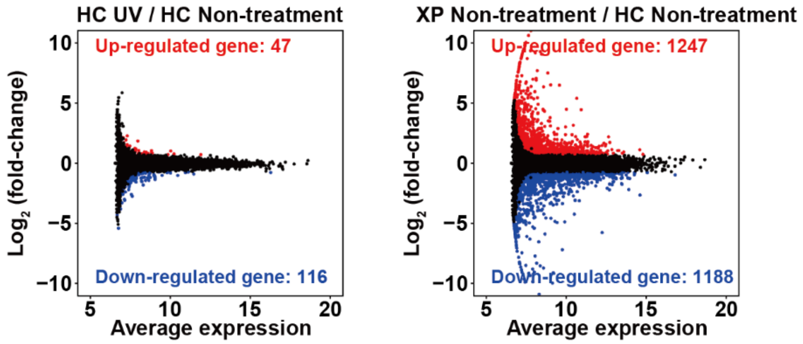


**Supplementary Figure 9. Gene expression alterations in HC-iMCs and XP-iMCs.**

The $\log_2$ (fold-change) in gene expression between HC-iMCs with and without UV irradiation (left) and between XP-iMCs and HC-iMCs without UV irradiation (right) was plotted against the average expression of each gene. DEGs were defined based on an FDR $< 0.05$ and $\log_2$ (fold-change) $> 0.7$ or $< -0.7$. Significantly up- and down-regulated genes are represented as red and blue dots, respectively.

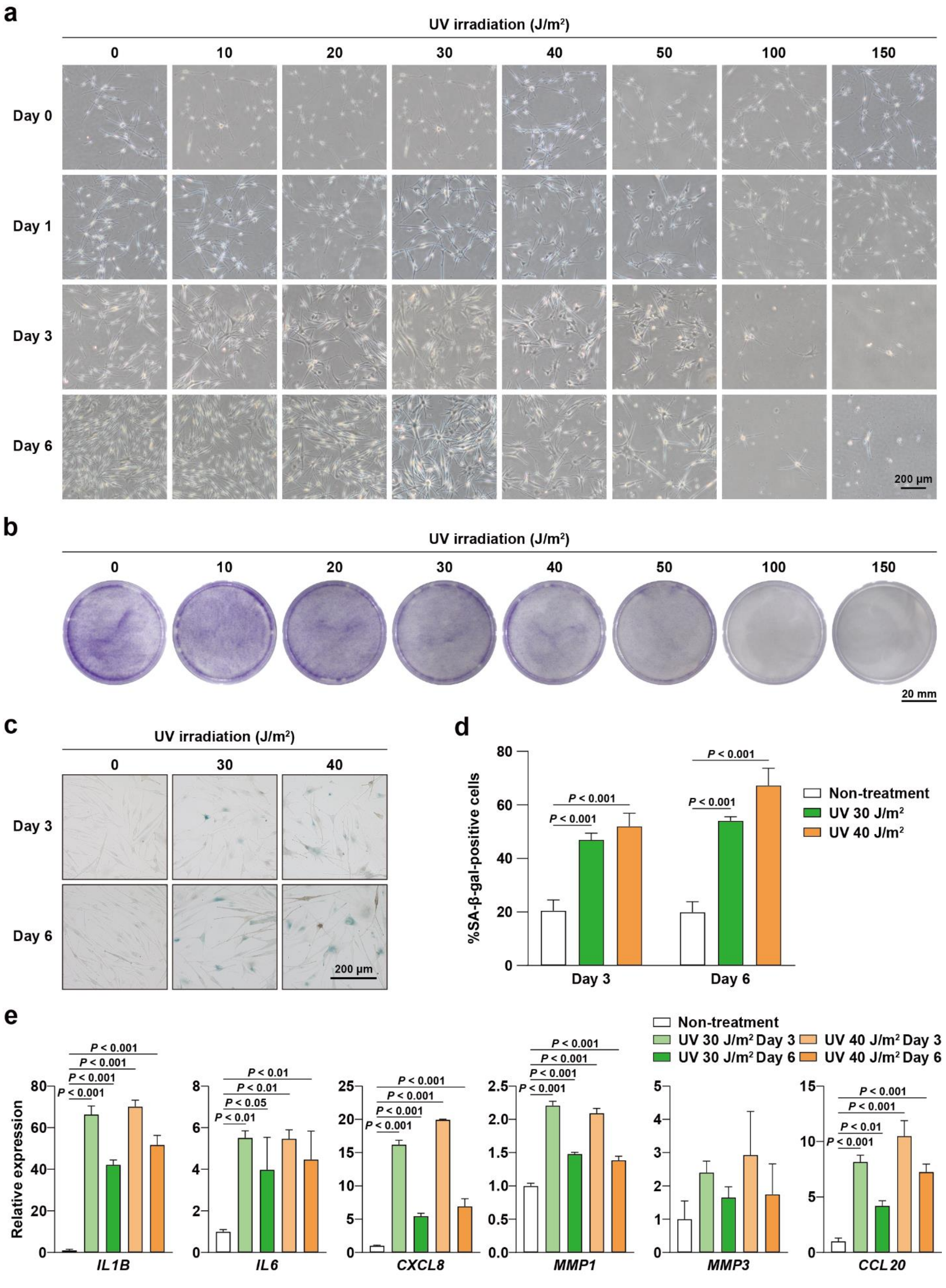
a
UV irradiation (J/m²)
0
10
20
30
40
50
100
150
Day 0
Day 1
Day 3
Day 6
200 µm
b
UV irradiation (J/m²)
0
10
20
30
40
50
100
150
20 mm
c
UV irradiation (J/m²)
0
30
40
Day 3
Day 6
200 µm
d
%SA-β-gal-positive cells
P < 0.001
Non-treatment
UV 30 J/m²
UV 40 J/m²
Day 3
Day 6
e
Non-treatment
UV 30 J/m² Day 3
UV 30 J/m² Day 6
UV 40 J/m² Day 3
UV 40 J/m² Day 6
Relative expression
P < 0.001
P < 0.01
P < 0.05
IL1B
IL6
CXCL8
MMP1
MMP3
CCL20

**Supplementary Figure 10. Effect of different UV doses on XP-iMCs.**

**(a)** Effect of different UV doses on XP-iMC morphology and viability. XP-iMCs are irradiated under the indicated UV conditions, and cell morphology images were captured at the specified time points (days).

**(b)** Effect of different UV doses on XP-iMC colony formation. Cells were irradiated under the indicated UV conditions, cultured for six days, and assessed for colony formation.

**(c)** SA-β-gal staining of XP-iMCs irradiated with UV under the specified conditions.

**(d)** Quantification of SA-β-gal-positive cells after UV irradiation. SA-β-gal staining images, as exemplified in **panel c**, were quantified.

**(e)** SASP gene expression in XP-iMCs irradiated with UV. Cells were irradiated under the indicated UV conditions, and gene expression was analyzed by RT-qPCR. Expression levels in non-irradiated XP-iMCs were set to 1.

In **panels d** and **e**, data represent means and standard deviations (n=3). Statistical significance was assessed using one-way ANOVA and Tukey's test.

#### Supplementary Table 1. List of senolytic agents

| Agents | Target | Pathway | Reference |
|---|---|---|---|
| ABT-263 | BCL-2 family protein | Apoptosis | (Chang et al., 2016; Wang et al., 2024; Zhu et al., 2016) |
| A-1331852 | BCL-X | Apoptosis | (Zhu et al., 2017) |
| A-1155463 | BCL-X | Apoptosis | (Zhu *et al.*, 2017) |
| S63845 | MCL-1 | Apoptosis | (Troiani et al., 2022) |
| BPTES | GLS1 | Glutaminolysis | (Johmura et al., 2021) |
| RSL3 | GPX4 | Ferroptosis | (Liao et al., 2022) |
| Digoxin | $Na^+/K^+$ ATPase pump | Concentration of $H^+$ | (Guerrero et al., 2019; Triana-Martinez et al., 2019) |
| Ouabain | $Na^+/K^+$ ATPase pump | Concentration of $H^+$ | (Guerrero *et al.*, 2019; Triana-Martinez *et al.*, 2019) |
| 17-DMAG | HSP90 | Apoptosis | (Fuhrmann-Stroissnigg et al., 2017) |
| Ganetespib | HSP90 | Apoptosis | (Fuhrmann-Stroissnigg *et al.*, 2017) |
| Panobinostat | Histone deacetylase | Gene repression | (Samaraweera et al., 2017) |
| Romidepsin | Histone deacetylase | Gene repression | (Bancaro et al., 2023) |

#### Supplementary Table 2. List of senomorphic agents

| Agents | Target | Pathway | Reference |
|---|---|---|---|
| A485 | Histone acetyltransferase p300 | Gene activation | (Sen et al., 2019) |
| Acetylcysteine | NF-κB | Inflammation | (Kunisada et al., 2017) |
| Curcumin | NF-κB | Inflammation | (Marquardt et al., 2015; Zia et al., 2021) |
| EF24 | NF-κB | Inflammation | (Li et al., 2019; Yin et al., 2016) |
| Metformin | NF-κB | Inflammation | (Moiseeva et al., 2013) |
| Apremilast | PDE4 | Gene repression | (Wang et al., 2021) |
| Difamilast | PDE4 | Gene repression | (Wang *et al.*, 2021) |
| Abrocitinib | JAK1 | JAK/STAT | (Xu et al., 2015) |
| Baricitinib | JAK1/2 | JAK/STAT | (Arnold et al., 2021; Liu et al., 2019; Xu *et al.*, 2015) |
| Upadacitinib | JAK1 | JAK/STAT | (Xu *et al.*, 2015) |
| IFN alpha-IFNAR-IN-1 | IFN-α/IFNAR | Type I interferon | (Wang *et al.*, 2024) |
| VBIT-4 | VDAC | Type I interferon | (Wang *et al.*, 2024) |
| Rapamycin | mTOR | mTOR | (Herranz et al., 2015) |
| Dasatinib | Tyrosine kinase | Apoptosis and Gene repression | (Wang et al., 2023; Zhu et al., 2015) |
| Quercetin | PI3K | Apoptosis and Gene repression | (Wang *et al.*, 2023; Zhu *et al.*, 2015) |

### Supplementary Table 3. List of IFN-I-related genes

| Description | Genes | Gene count | IFN- I genes | IFN- I gene count |
|---|---|---|---|---|
| ***Figure 7d*** | | | | |
| Cytokine-cytokine receptor interaction | *CXCR4, TNFRSF18, CXCL8, CCL20, LIF, NGF, IL17F, CCL25, LEP, IL12B, IFNA5, TSLP, CXCL14, EPO, IL5RA, AMHR2, OSM, CD27, TNFRSF10C, IL10RA, TNFRSF1B, CSF3, IL13, IL15RA, IL12A, LTA, IL1B, BMP8B, CLCF1, GDF6, TNFRSF9, IL19, CCR7, INHBA, CD70, GDF5, TNFRSF4, CXCL17, IL1RL2, TNFRSF10D, IL18RAP, CCR4, IL10, CXCL3, TNFSF9, CNTFR, IL31RA, IL17C, IL21R, TGFB3, INHBB, IFNA1, CCL26, XCR1, IL22RA1, TNFRSF13C, INHBC, IL11, CD4, TNFRSF12A, AMH, THPO, IL17RC, IL2RA, BMP2, TNFSF10, TNFRSF17, IL2RG, TNFRSF10A, TNFRSF11A, TNFSF13B, CCL17, IL5, IL34, IL23A, IL17B, IL13RA2, MPL, CXCL6, CCL28, BMP5, IL37, IL31, IL20, IL18, IL7R, CXCL2, IL32, IL18R1, FAS, NODAL, IL6, CCL24, PRLR, CCL22, TNFRSF8, CXCL1* | 97 | *IFNA5, IFNA1* | 2 |
| Herpes simplex virus 1 infection | *CFP, C3, IL12B, P3R3URF-PIK3R3, ZNF560, IFNA5, ZNF705A, CD74, IL12A, LTA, IL1B, ZNF547, ZNF773, BIRC3, ZNF559-ZNF177, ZNF596, IRF7, ZIM3, IFNA1, ZNF454, ZNF568, ZNF707, ZNF699, ZNF10, ZNF559, ZNF597, HLA-DMB, ZNF425, ZNF721, NXF1, ZNF732, ZNF19, ZNF211, ZNF28, ZNF614, ZNF436, ZNF669, ZNF460, SRSF1, ZNF415, ZNF426, ZNF836, ZNF439, POU2F2, ZNF556, ZNF585B, FAS, ZNF570, ZNF544, ZNF442, ZNF14, ZNF584, ZNF253, ZNF525, ZIK1, OAS2, ZNF785, ZNF600, ZNF514, IL6, ZNF805, ZNF517, ZNF778, ZNF432, ZNF350, ZNF888, ZNF92, ZNF264, HLA-B, ZNF470, ZNF615, ZNF845, ZNF540, ZNF256, ZNF583, MYD88, ZNF124, ZNF808, ZNF571, ZNF667, ZNF850, ZNF225, ZNF530, ZNF573, ZNF607, ZNF557, ZNF317, ZNF416, ZNF582, ZNF543, ZNF701, ZNF331, ZNF79, ZNF846, ZNF304, ZNF783, ZNF534, PILRA, ZNF497, ZNF829, ZNF620, ZSCAN32, ZNF473, ZNF20, ZNF77, ZNF586, ZNF611, ZNF606, ZNF461, HLA-C, ZNF550, ZNF136, ZNF383, ZNF418, ZNF552, ZNF195, ZNF26, ZNF98, IFNAR2, ZNF728, ZNF311, ZNF181, ZNF577, ZNF677, ZNF227, ZNF12, ZNF549, ZFP69B, ZNF251, ZNF441, OAS3, ZNF200, ZNF224, ZNF649, ZNF880, BST2, ZNF490, ZNF765, ZNF343, EIF2B2, ZNF529, ZNF891, ZNF791, ZNF101, ZNF267, ZNF234, ZNF761, ZNF610, ZNF510, ZNF689, ZNF567, ZNF302, ZNF155, ZNF3, ZNF160, ZNF561, ZNF284, ZNF738, RELA, ZNF814, POU2F3, HLA-E, ZNF324, ZNF558, ZNF623, B2M, ZNF554, ZNF726, ZFP1, ZNF682, BIRC2, ZNF621, TAP1, ZNF684, ZNF141, ZNF84, ZNF175, TRAF6, ZNF587, ZNF226, EIF2S1, ZNF268, CGAS, ZFP37, ZNF768, ZNF337, ZNF235, ZNF527, ZNF484, ZNF528, ZNF2, ZNF563, TP53, ZNF140, ZNF354B, ZNF404, ZNF485, ZNF766, ZNF7, ZNF718, ZNF786, ZNF180, CYCS, CASP3, ZNF724, ZNF671, ZNF790, EIF2B1, ZNF519, ZNF250, ZNF813, ZNF354C, ZNF569, SOCS3, ZNF675, ZNF776, ZNF320, SRSF8, ZNF93, SRSF6, ZNF266, ZNF205, BAK1* | 223 | *IFNA5, IRF7, IFNA1, OAS2, MYD88, IFNAR2, OAS3* | 7 |
| JAK/STAT signaling pathway | *LIF, LEP, IL12B, PTPN6, P3R3URF-PIK3R3, IFNA5, TSLP, EPO, IL5RA, OSM, IL10RA, CSF3, IL13, IL15RA, IL12A, CLCF1, PDGFB, IL19, IL10, SOCS1, CNTFR, IL31RA, IL21R, AOX1, IFNA1, IL22RA1, IL11, THPO, IL2RA, CCND2, IL2RG, IL5, CDKN1A, IL23A, IL13RA2, MPL, IL31, IL20, IL7R, CISH, IL6, PRLR, SOCS4, IL15, PDGFRB, CSF3R, LIFR, IL23R, IFNAR2* | 49 | *PTPN6, IFNA5, IFNA1, IFNAR2* | 4 |
| ***Figure 7g*** | | | | |
| Human papillomavirus infection | *THBS1, MDM2, CDKN1A, FAS, ITGA7, FZD4, HES2, PDGFRB, ISG15, MX1, CREB5, VEGFA, COL9A3, CREB3L1, FOXO1, WNT8B* | 16 | *ISG15* | 1 |
| MAPK signaling pathway | *NGFR, GDNF, FAS, GADD45A, PLA2G4C, RASGRF1, KITLG, PGF, PDGFRB, PDGFC, VEGFA, DUSP5, MYD88, AREG, FGFR3* | 15 | *MYD88* | 1 |
| Influenza A | *TNFRSF10B, FAS, OAS3, IRF7, OAS2, TNFRSF10A, MX1, CXCL8, MYD88* | 9 | *OAS3, IRF7, OAS2, MYD88* | 4 |
| ***Supplementary Figure 7b*** | | | | |
| Influenza A | *KPNA1, OAS1, TNFRSF10B, FAS, IL33, CASP1, OAS3, IRF7, OAS2, STAT1, TNFRSF10A, MX1, TRIM25, TLR3, ADAR, IL1B, CXCL8, IFIH1, EIF2AK2, MYD88, RSAD2* | 21 | *OAS1, OAS3, IRF7, OAS2, STAT1, ADAR, IFIH1, MYD88* | 8 |
| Human papillomavirus | *THBS1, FN1, MDM2, LAMA1, CCND1, ITGA3, CDKN1A, SPP1, LAMB1, NOTCH3, FAS, TNC, THBS2, ITGA1, ITGA7, FZD4,* | 38 | *ISG15, STAT1* | 2 |

| | | | | |
|---|---|---|---|---|
| infection | *ITGA4, HEY1, HES2, PDGFRB, LAMB3, ISG15, STAT1, MX1, CREB5, VEGFA, PATJ, COL9A3, TLR3, LAMA2, OASL, CREB3L1, FOXO1, EIF2AK2, THBS4, ATP6V0D2, HEYL, WNT8B* | | | |
| Measles | *CCND1, IRAK1, OAS1, FAS, TNFAIP3, OAS3, IRF7, OAS2, STAT1, MX1, ADAR, IL1B, IFIH1, EIF2AK2, MYD88* | 15 | *IRAK1, OAS1, OAS3, IRF7, OAS2, STAT1, ADAR, IFIH1, MYD88* | 9 |
| Malaria | *THBS1, THBS2, PECAM1, IL1B, CXCL8, THBS4, MYD88, ITGB2* | 8 | *MYD88* | 1 |
| AGE-RAGE signaling pathway in diabetic complications | *FN1, CCND1, SERPINE1, THBD, EGR1, STAT1, VEGFA, IL1B, PLCB2, CXCL8, VEGFC, FOXO1, PLCE1* | 13 | *STAT1* | 1 |
| Viral life cycle - HIV-1 | *MAP1A, TRIM5, SAMHD1, MX1, MAP1B, APOBEC3G, BST2, APOBEC3H, CD4* | 9 | *SAMHD1* | 1 |
| Tuberculosis | *ITGAX, IRAK1, SPHK1, PLK3, PLA2R1, VDR, STAT1, IL1B, IRAK2, ATP6V0D2, CORO1A, CTSS, TLR1, MYD88, IL23A, ITGB2* | 16 | *IRAK1, STAT1, MYD88* | 3 |
| ***Supplementary Figure 7d*** | | | | |
| Cytokine-cytokine receptor interaction | *CCR1, IL1B, CXCL8, CXCL6, IL31RA, LIF, IL2RB, CCR3, IL12A, IL7R, NODAL, IFNA5, NGFR, IL1RL2, OSM, TNFSF13B, CSF2RB, CCR7, IFNB1, NGF, CCL24, LEP, IL11, EBI3, IL33, BMP8B, IL5, CCR4, CXCR3, MPL, TNFRSF11A, CXCL1, TNFRSF8, CCL3L1, CCL25, TNFSF11, CXCR4, IL17C, CNTF, INHBB, CCL28, IFNLR1, TSLP, IL18R1, TNFRSF17, TNFRSF12A, IL4, IL23A, TNFRSF1B, CXCR6, TNFRSF10A, IL12RB2, INHBA, TNFSF9, TNFRSF10C, CCL27, IL1R1, GDF6, ACVRL1, TNFRSF21, CD40LG, IL13RA2, CSF3R, TNFSF15, IL20, IL6, IL27, IL22RA1, TNFRSF13B, IL20RA, CSF3, IL12B, CXCL5, CXCL17, LIFR, FAS, GDF15, BMP7, CCL3, BMPR1B, CD4, CD27, CSF1, CXCL12, TNFRSF19, TNFRSF10B, CLCF1, CNTFR, IL4R, IL1RAP, GDF5, PRL* | 92 | *IFNA5, IFNB1* | 2 |
| Influenza A | *IL1B, CXCL8, OAS2, OAS1, IL12A, IFNA5, HLA-DOA, MX1, OAS3, IFNB1, KPNA7, IL33, RSAD2, IRF7, HLA-DMB, HLA-DQB1, HLA-DQB2, MX2, IFIH1, TNFRSF10A, STAT1, EIF2AK2, NXF3, IL6, IL12B, FAS, TLR3, TRIM25, CASP1, KPNA1, MYD88, ADAR, TNFRSF10B* | 33 | *OAS2, OAS1, IFNA5, OAS3, IFNB1, IRF7, IFIH1, STAT1, MYD88, ADAR* | 10 |
| JAK/STAT signaling pathway | *PDGFB, IL31RA, LIF, IL2RB, IL12A, IL7R, IFNA5, OSM, PDGFRB, CSF2RB, IFNB1, LEP, PDGFRA, IL11, IL5, MPL, CNTF, IFNLR1, TSLP, AOX1, IL4, IL23A, STAT4, IL12RB2, STAT1, EGF, IL13RA2, CSF3R, IL20, IL6, IL27, IL22RA1, IL20RA, CSF3, IL12B, CCND1, LIFR, CDKN1A* | 38 | *IFNA5, IFNB1, STAT1* | 3 |
| Tuberculosis | *IL1B, CORO1A, ATP6V0D2, NOS2, ITGAX, IL12A, IFNA5, CD14, CR1L, HLA-DOA, NOD2, IFNB1, CASP10, HLA-DMB, HLA-DQB1, HLA-DQB2, CAMP, CTSD, CTSS, IL23A, IRAK2, CAMK2A, VDR, LSP1, FCGR1A, STAT1, CYP27B1, ITGB2, C3, CR1, IL6, IL12B, PLK3, SPHK1, PLA2R1, IRAK1, TLR1, MYD88* | 38 | *IFNA5, IFNB1, STAT1, IRAK1, MYD88* | 5 |
| Human papillomavirus infection | *HES2, ATP6V0D2, WNT8B, WNT9A, IFNA5, LAMC2, MX1, PDGFRB, OASL, THBS1, IFNB1, ISG15, WNT3A, MFNG, LAMA1, NOTCH4, WNT9B, HES3, ITGB6, RBPJL, TNC, PATJ, TBPL2, MX2, LAMB3, CREB5, TNXB, ITGA10, THBS2, MDM2, HEYL, HEY1, ITGA4, FN1, COL4A6, WNT10B, CREB3L1, WNT3, SPP1, STAT1, ITGA3, EIF2AK2, LAMA2, EGF, ITGA1, HES5, NOTCH3, WNT7A, CCND1, FZD4, CDKN1A, ITGA7, FOXO1, FAS, TLR3, ITGA8, WNT6, ITGB3, COL9A3, COL9A2, LAMB1, THBS4, VEGFA, IRF1, LAMA3, ITGA5, ITGB1, FZD7, NOTCH1, COL1A2, COL4A1, ITGA6, ITGA2* | 73 | *IFNA5, IFNB1, ISG15, STAT1* | 4 |
| ***Supplementary Figure 8d*** | | | | |
| Measles | *OAS1, OAS3, IRF7, OAS2, STAT1, MX1, EIF2AK2, STAT2, PIK3R1* | 9 | *OAS1, OAS3, IRF7, OAS2, STAT1, STAT2* | 6 |
| Hepatitis C | *OAS1, IFIT1, OAS3, IRF7, OAS2, STAT1, MX1, EIF2AK2, STAT2, PIK3R1* | 10 | *OAS1, OAS3, IRF7, OAS2, STAT1, STAT2* | 6 |
| Influenza A | *OAS1, OAS3, IRF7, OAS2, STAT1, MX1, EIF2AK2, STAT2, PIK3R1* | 9 | *OAS1, OAS3, IRF7, OAS2, STAT1, STAT2* | 6 |
| Epstein-Barr virus infection | *MDM2, OAS1, OAS3, IRF7, OAS2, ISG15, STAT1, EIF2AK2, STAT2, PIK3R1* | 10 | *OAS1, OAS3, IRF7, OAS2, ISG15, STAT1, STAT2* | 7 |
| NOD-like receptor signaling pathway | *OAS1, OAS3, IRF7, OAS2, STAT1, STAT2, NAMPT, IFI16, ERBIN* | 9 | *OAS1, OAS3, IRF7, OAS2, STAT1, STAT2* | 6 |

| | | | | |
|---|---|---|---|---|
| Coronavirus disease - COVID-19 | *OAS1, OAS3, OAS2, ISG15, STAT1, MX1, EIF2AK2, STAT2, PIK3R1, IL6ST* | 10 | *OAS1, OAS3, OAS2, ISG15, STAT1, STAT2* | 6 |
| Human papillomavirus infection | *THBS1, FN1, MDM2, ITGA1, ISG15, STAT1, MX1, EIF2AK2, STAT2, PIK3R1, FZD3, MAML2* | 12 | *ISG15, STAT1, STAT2* | 3 |
| Viral carcinogenesis | *MDM2, HDAC9, IRF7, SP100, EIF2AK2, PIK3R1, IL6ST* | 7 | *IRF7, SP100* | 2 |
| ***Supplementary Figure 8e*** | | | | |
| Hepatitis C | *EGF, CLDN1, TLR3, CLDN20, CLDN5, STAT1, SOCS3, EIF2AK2, E2F2, CLDN16, CLDN19, P3R3URF-PIK3R3, CLDN14, CLDN12, CLDN2, MX2, IRF7, OAS3, MX1, IFNB1, IFNA5, OAS1, IFIT1, RSAD2, OAS2* | 25 | *STAT1, IRF7, OAS3, IFNB1, IFNA5, OAS1, OAS2* | 7 |
| Influenza A | *TMPRSS11D, TRIM25, TLR3, IL33, HLA-DMB, STAT1, SOCS3, IL12A, IFIH1, EIF2AK2, IL12B, IL1B, P3R3URF-PIK3R3, IL1A, MX2, HLA-DQB2, HLA-DQB1, IRF7, OAS3, MX1, IFNB1, HLA-DOA, IFNA5, OAS1, RSAD2, OAS2* | 26 | *STAT1, IFIH1, IRF7, OAS3, IFNB1, IFNA5, OAS1, OAS2* | 8 |
| Epstein-Barr virus infection | *PLCG2, CIR1, HLA-B, HLA-DMB, STAT1, EIF2AK2, E2F2, ITGAL, P3R3URF-PIK3R3, BLNK, CD3G, ENTPD8, HLA-DQB2, HLA-DQB1, IRF7, OAS3, IFNB1, ISG15, HLA-DOA, IFNA5, OAS1, OAS2* | 22 | *STAT1, IRF7, OAS3, IFNB1, ISG15, IFNA5, OAS1, OAS2* | 8 |
| Coronavirus disease - COVID-19 | *STAT1, IL12A, IFIH1, EIF2AK2, IL12B, MBL2, CSF3, MASP1, RPL3L, IL1B, P3R3URF-PIK3R3, C7, MMP3, MX2, C5AR1, OAS3, MX1, RPL17-C18orf32, IFNB1, ISG15, IFNA5, PRKCG, OAS1, OAS2* | 24 | *STAT1, IFIH1, OAS3, IFNB1, ISG15, IFNA5, OAS1, OAS2* | 8 |
| JAK/STAT signaling pathway | *PDGFB, MPL, CLCF1, PDGFRB, IL23R, IL4, IFNK, IL23A, STAT1, SOCS3, IL12A, IL12B, CSF3, IL20RA, IL22RA1, EPO, P3R3URF-PIK3R3, PDGFRA, IL7R, IL2RB, IL2RA, PRL, IL31RA, LEP, IFNB1, CSF2RB, OSM, TSLP, IFNA5* | 29 | *IFNK, STAT1, IFNB1, IFNA5* | 4 |
